# Oscillations and variability in neuronal systems: interplay of autonomous transient dynamics and fast deterministic fluctuations

**DOI:** 10.1101/2021.06.14.448371

**Authors:** Rodrigo F. O. Pena, Horacio G. Rotstein

**Author notes:** Corresponding Investigator, CONICET, Argentina.

## Abstract

Neuronal systems are subject to rapid fluctuations both intrinsically and externally. These fluctuations can be disruptive or constructive. We investigate the dynamic mechanisms underlying the interactions between rapidly fluctuating signals and the intrinsic properties of the target cells to produce variable and/or coherent responses. We use linearized and non-linear conductance-based models and piecewise constant (PWC) inputs with short duration pieces. The amplitude distributions of the constant pieces consist of arbitrary permutations of a baseline PWC function. In each trial within a given protocol we use one of these permutations and each protocol consists of a subset of all possible permutations, which is the only source of uncertainty in the protocol. We show that sustained oscillatory behavior can be generated in response to various forms of PWC inputs independently of whether the stable equilibria of the corresponding unperturbed systems are foci or nodes. The oscillatory voltage responses are amplified by the model nonlinearities and attenuated for conductance-based PWC inputs as compared to current-based PWC inputs, consistent with previous theoretical and experimental work. In addition, the voltage responses to PWC inputs exhibited variability across trials, which is reminiscent of the variability generated by stochastic noise (e.g., Gaussian white noise). Our analysis demonstrates that both oscillations and variability are the result of the interaction between the PWC input and the target cell’s autonomous transient dynamics with little to no contribution from the dynamics in vicinities of the steady-state, and do not require input stochasticity.

## 1 Introduction

Variability in neuronal activity has been observed at all levels of neuronal organization [1–14]. However, the mechanisms underlying the generation of variable voltage responses to fluctuating inputs are not well understood. In particular, it is not well understood which aspects of the variable voltage responses of neuronal systems are due to their intrinsic properties (e.g., ionic currents), which aspects are due to properties of the external inputs, and how the two interact.

The goal of this paper is to address these issues in the context of single cells receiving external current inputs. Through a series of case studies, we systematically investigate the role of the intrinsic properties of the target cells in the generation of oscillatory and variable responses to fluctuating inputs, and how the response variability is controlled by these intrinsic properties.

The cellular intrinsic and dynamic properties can be uncovered and characterized by the use of constant inputs. The voltage responses to constant input currents consist of a transient phase followed by a stationary phase [15]. Examples of transient behavior are monotonic changes towards equilibrium, overshoots (or sags), and damped oscillations. We collectively refer to them as the autonomous transient dynamics. These two phases reflect different ways in which the cellular intrinsic properties interact with the input and therefore different ways in which the voltage response encodes information about these cellular intrinsic properties. We argue that the autonomous transient dynamics plays a key role in determining the cell’s variable response to external inputs, while the cell’s stationary response to constant inputs plays at most a minor role. We additionally show that variability emerges in response to multiple presentations of (deterministic) piecewise constant (PWC) input functions having exactly the same constant pieces arranged in different, arbitrary orders across trials. This provides a way of elucidating the mechanistic role of both the cellular intrinsic properties through the corresponding autonomous transient dynamics in generating the response variability. This also supports the idea that variability, although it is present in response to stochastic inputs, is not inherently a stochastic phenomenon.

A prototypical example of the role of the transient dynamics of individual cells (autonomous transient dynamics) in determining their response to external inputs is given by subthreshold resonance [16–18]. Subthreshold resonance refers to the ability of cells to exhibit a peak in the impedance amplitude profile (curve of the impedance amplitude as a function of the input frequency, Figs. 2-b3 and -c3) for a preferred (resonant) input frequency [19, 20] (see Appendix A). Subthreshold resonance emerges in both cells having intrinsic oscillatory properties (e.g., two-dimensional linear cells with complex eigenvalues, Fig. 2-c3) and cells lacking intrinsic oscillatory dynamics [16,21] (e.g., two-dimensional linear cells having real eigenvalues, Fig. 2-b3), but exhibiting an overshoot (or sag). The effective time scale governing these autonomous transient dynamics is key for the generation of subthreshold resonance and the determination of the resonant frequency, which reflects balances between interacting processes. [16–18].

Neurons and neuronal networks are subject to intrinsic random fluctuations [22–24] such as random opening/closing of ionic channels [25–32] and random synaptic inputs from other neurons in the network [1, 33–38]. Random fluctuations are also exerted by external factors. In mathematical models, these fluctuations are incorporated as noise interacting in various ways with the underlying deterministic dynamics [39].

Noise plays many roles in the resulting patterns. It can uncover the transient dynamics of dynamical systems [40]. It can also modify the dynamics prescribed by the underlying deterministic system [41], for example, by creating sustained (irregular) oscillations in systems that would exhibit an equilibrium otherwise [42] or by creating bursting (spiking) patterns in systems that would exhibit deterministic spiking (bursting) patterns otherwise. From one point of view, noise is disruptive in the sense that it creates irregularities in otherwise regular patterns or even destroys equilibrium patterns. From another perspective, noise is constructive in the sense that it may create oscillatory patterns or increase the signal coherence. Two prototypical examples are stochastic resonance [43–51] and coherence resonance [52–59]. Additionally, noise can induce order in chaotic dynamics [60], promote synchronization (stochastic synchronization) [61–63] or generate synchronized oscillations in networks of coupled excitable elements [64–67], promote the generation of slow wave oscillations by inducing transitions between active and silente phases [68, 69], and induce transitions in bistable and excitable systems (non-oscillatory) [29, 58, 70–74] among other phenomena.

Common to these phenomena is the idea that noise repeatedly “kicks” trajectories and keeps them away from their stationary solutions (when they exist) to create alternative patterns (irregular versions of the underlying noiseless pattern or qualitatively different patterns). In practice, external noise interacts with the cell, which responds by integrating the intrinsic (deterministic) and input (stochastic) components (similarly to what was described above). A prototypical example of a stochastic input to a neuron is the Ornstein-Uhlenbeck (OU) process [75–77] (Appendix B). While the response’s expected value coincides with the stationary solution of the noiseless system, the response’s variance involves a combination of the (noiseless) model parameters and the variance of the Gaussian white noise input. In other words, the response variability is controlled by the autonomous transient dynamics. For higher-dimensional OU process, the determination of the dependence of the variability with the autonomous transient dynamics is more difficult than for one-dimensional OU processes given the complexity of the covariance formulas. This type of determination is analytically not possible for noise-driven nonlinear systems (including voltage-dependent nonlinearities or conductance-based synaptic inputs). These tasks require a different approach and the development of a conceptual framework that allows the investigation of the response properties in terms of the properties of the participating building blocks: the target cells and the fluctuating inputs. The autonomous transient dynamics are a key element of this framework.

Here, we use piecewise constant (PWC) input currents (*I_η_*) with short duration pieces and variable amplitudes that allow for the autonomous transient dynamics to develop in response to each constant piece input. When the set of amplitudes (*η*) are normally distributed and the durations are small enough, *I_η_* is an approximation to Gaussian white noise [78]. However, in this paper we are not interested in this limit. We first show that an additive input *I_η_* with normally distributed amplitudes evoke oscillatory voltage responses in cells whose stable equilibria are either foci or nodes (displaying damped oscillations and overshoots in response to constant inputs, respectively). We note that the former captures results described in [76] for Gaussian white noise (see also [79] for two-dimensional linear cells with damped oscillations). We then use the same input to investigate the response properties of a nonlinear (piecewise linear, PWL) model [17], which mimics the voltage-dependencies present in neurons, and a linear model receiving a (multiplicative) conductance-based synaptic-like current. We explain how oscillations are nonlinearly amplified or attenuated in these models. Next, we investigate how the voltage response variability is linked to the properties of the autonomous transient dynamics. All trials for a given protocol have the same set of linear piece amplitudes *η*, but the order of the constant pieces is different for each trial. Each protocol consists of a subset of all possible order permutations of the elements of the set *η*. This is the only source of uncertainty in the process. We compute the peak-and-troughs voltage response profiles *P_η_* consisting of the set of peaks and troughs of the voltage response evoked by each constant piece, arranged in an appropriate way for comparison. We analyze the dependence of the variability of the *P_η_* patterns across trials with the properties of the autonomous transient dynamics, and we use these results to understand how the response variability depends on the model parameters. More specifically, this variability results from the multiple different ways a target cell reacts to the same constant inputs (across trials) as it transitions from the response to the previous constant piece. Finally, we show how frequency-dependent inputs affect these processes.

## 2 Methods

### 2.1 Models

In this paper we use relatively simple biophysically plausible models describing the subthreshold dynamics of individual neurons subject to both additive and multiplicative inputs.

#### 2.1.1 Linear model: additive input

For the individual neurons we use the following linearized biophysical (conductance-based) model [16, 21]

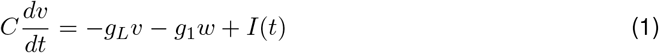

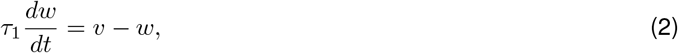

where *v* (mV) is the membrane potential relative to the voltage coordinate of the fixed-point (equilibrium potential) of the original model, *w* (mV) is the recovery (gating) variable relative to the gating variable coordinate of the fixed-point of the original model normalized by the derivative of the corresponding activation curve, *C* (*μ*F/cm^2^) is the specific membrane capacitance, *g_L_* (mS/cm^2^) is the linearized leak conductance, *g*_1_ (mS/cm^2^) is the linearized ionic conductance, *τ*_1_ (ms) is the linearized gating variable time constant and *I*(*t*) (*μ*A/cm^2^) is the time-dependent input current. In this paper we consider resonant gating variables (*g*_1_ > 0; providing a negative feedback effect). The linearization process for conductance-based models for single cells has been previously described in [16, 21]. We refer the reader to these references for details.

#### 2.1.2 Piecewise linear (PWL) model: additive input

To account for nonlinear effects we extend the linear model (1)-(2) to include a piecewise linear function *F_P W L_* [17]

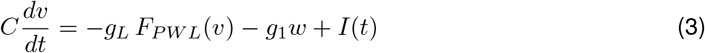

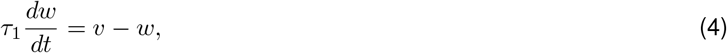

where

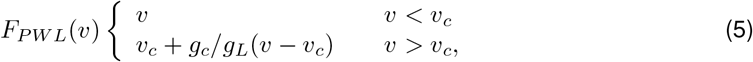

where *v_c_* is the cutting point (or breaking point) of the PWL model. This model has been used to investigate the dynamic mechanisms underlying the nonlinear amplification of the resonant voltage responses to sinusoidal inputs [17] and captures the nonlinear amplification effects of the resonant voltage responses off two-dimensional models with parabolic-like voltage nullclines [18]. Note that the PWL model has linear dynamics for *v < v_c_* and therefore the PWL model becomes effectively linear if *v_c_* is large enough.

#### 2.1.3 Conductance-based synaptic input model: multiplicative input

To account for the effects of conductance-based synaptic inputs we extend the model (3)-(5) to include a synaptic current

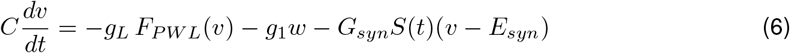

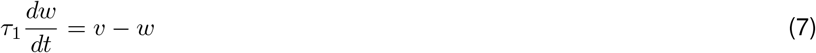

where *G_syn_* (mS/cm^2^) is the maximal synaptic conductance, *E_syn_* (mV) is the synaptic reversal potential and *S*(*t*) is the time-dependent synaptic input. We use *E_syn_* = 1 (excitatory synaptic current).

### 2.2 Piecewise constant (PWC) input functions with variable amplitudes

We use PWC functions with short-duration pieces as a tool to understand how the properties of the transient voltage response of individual cells to constant inputs (autonomous transient dynamics) control the response variability of these cells to time-dependent inputs with variable, abruptly changing amplitudes. White noise and related stochastic inputs (e.g. colored noise) satisfies these last two properties. PWC inputs with variable amplitudes and short durations provide a good balance between input variability and the ability of the autonomous transient dynamics to develop, and therefore they can be linked to the input amplitude that gave rise to them. In addition, PWC inputs with variable amplitudes allow us to use the same set of amplitudes in all trials for each protocol, arranged in different order for each trial, and therefore have a better control of the process. PWC functions with normally distributed amplitudes have been used to approximate white noise [78] (see also [80]) and converge to white noise as the duration of all pieces approaches zero.

#### 2.2.1 Piecewise constant input functions with normally distributed amplitudes

We partition the time interval [0, *T_max_*] into *N* pieces of equal length Δ, and we define *I_η_*(*t*) = *η_k_* for *t* ∈ (*t_*K* − 1_, t_k_*) for *k* = 1, ... *N*. The values of the amplitudes 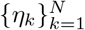 of the constant pieces are normally distributed (zero mean and variance *D*) (Fig. 1, top, see also Fig. 3-a). For each protocol in our study we use multiple different input trial functions *I_η_*(*t*) constrained to consist of different (random) permutations of the same set 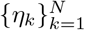. As part of our analysis we use one specific rearranged version *I_η,step_* of *I_η_* where the linear pieces 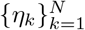 are ordered in a monotonically increasing amplitude manner (Fig. 1, middle). We use this as our notion of a reference (“ordered”) input in the sense that *I_η,step_* has the minimum piece-to-piece variability (except possibly for the decreasing order).

**Figure 1:**
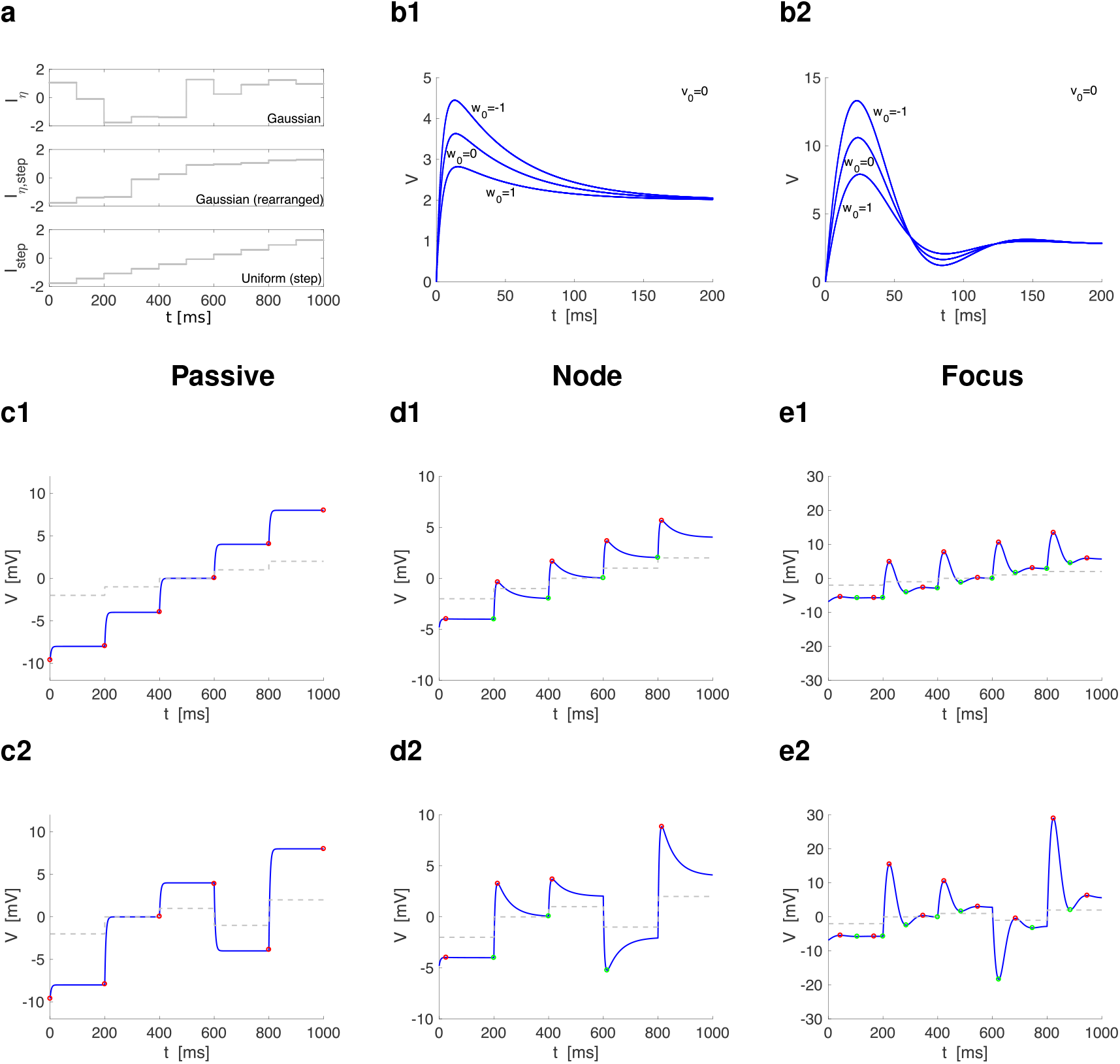
Representative voltage responses of linear systems to piecewise constant inputs with normally distributed amplitudes. **a.** Representative examples of piecewise constant inputs (*N* = 10, Δ = 100). Top: the values of the constant pieces of *I_η_* are normally distributed (mean zero and variance one). Middle: the values of the constant pieces of *I_η,step_* are as in the top panel, but rearranged in increasing order. Botton: the values of the constant pieces of *I_step_* are such that the steps are equal with *η*_1_ and *η_N_* taken from *I_η,step_* (bottom). In other words, *I_η,step_* and *I_step_* change in between the same values (*η*_1_ and *η_N_*) across trials. **b.** Dependence of the overshoot (b1) and first oscillation (b2) peaks with the initial value *w*(0) = *w*_0_ for the recovery variable *w* in response to the same constant input *I_app_* = 1 and *v*(0) = 0. Changes in the initial value of *v* produces only minor changes in the response patterns. **c to e.** Response of cells with three qualitatively different types of autonomous dynamics to piecewise constant inputs (*N* = 5, Δ = 200). Red and green dots indicate peaks and troughs, respectively. **Row 2.** *I_step_* ordered by increasing values of 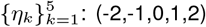. **Row 3.** *I_η_* using the values of 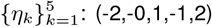 as in row 2, but with a different (non-monotonic) distribution. **c.** Passive cell. We used the following parameter values: *C* = 1, *g_L_* = 0.25. Red dots indicate the boundaries between constant pieces (there are no peaks and troughs). **d.** 2D linear system exhibiting an overshoot in response to step-constant inputs. We used the following parameter values: *C* = 1, *g_L_* = 0.25, *g*_1_ = 0.25, *τ*_1_ = 100. **e.** 2D linear system exhibiting damped oscillations in response to step-constant inputs. We used the following parameter values: *C* = 1, *g_L_* = 0.05, *g*_1_ = 0.3, *τ*_1_ = 100.

**Figure 2:**
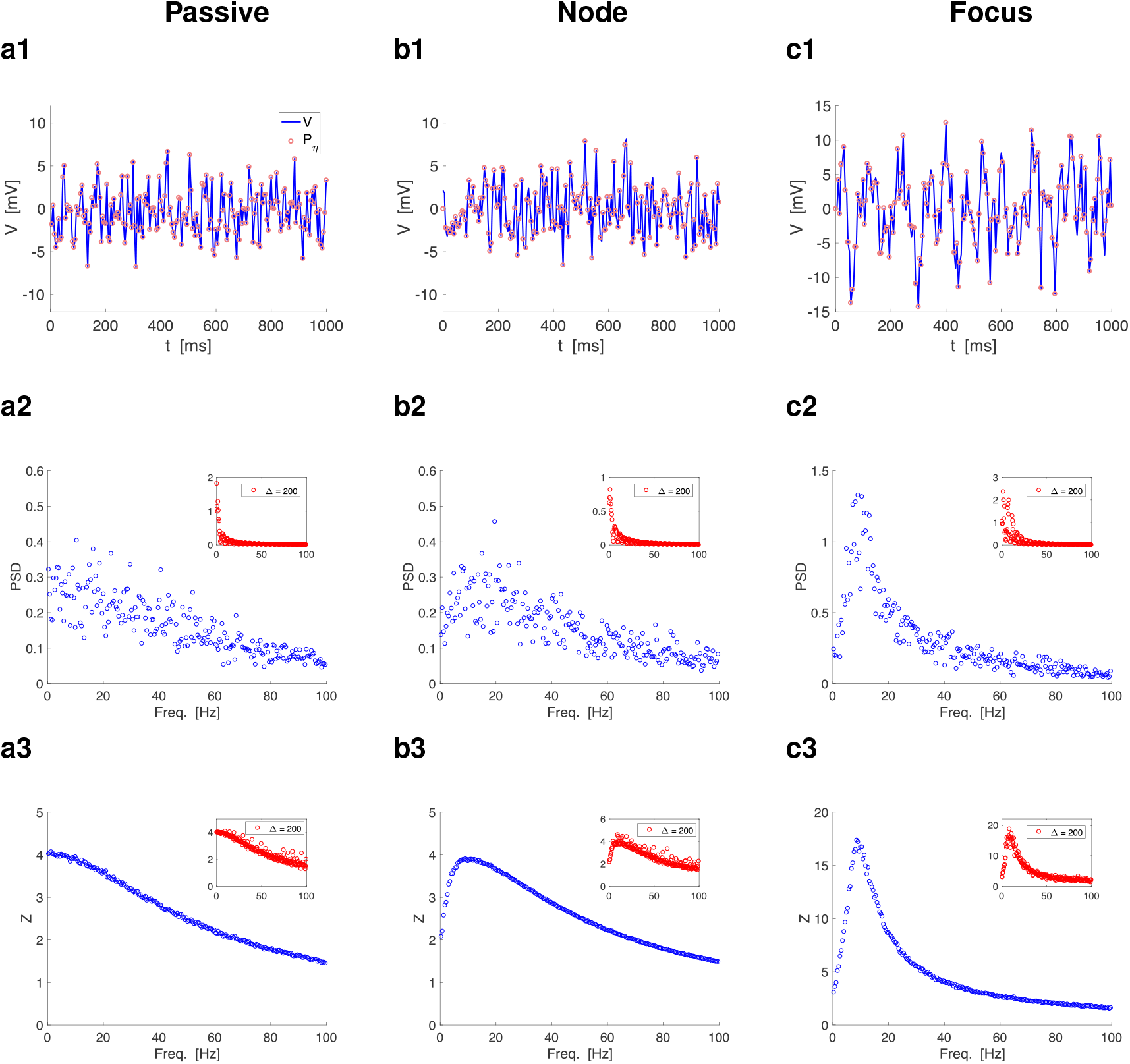
Piecewise constant inputs with normally distributed amplitudes capture the transient dynamics of the target cells. The piecewise constant inputs *I_η_* have Δ = 5 (total time = 10000 ms, N = 2000). The insets have Δ = 200 (total time = 10000 ms, N = 50). The parameter values are the same as in Fig. 1. **Row 1.** Representative *V* traces (only 1000 ms are shown). The coral dots indicate peaks-and-troughs patterns computed as the maximum (minimum) of the *V* response if *η_k_* > η_*k* − 1_ (*η_k_* < η_*k* − 1_). **Row 2.** Power spectra density (PSD) profiles for the sample *V* trace. **Row 3.** Impedance amplitude (*Z*) profiles for the sample *V* trace. **a.** Passive cell (*f_nat_* = *f_res_* = 0). We used the following parameter values: *C* = 1, *g_L_* = 0.25. **b.** 2D linear system exhibiting an overshoot in response to step-constant inputs ((*f_nat_* = 0*, f_res_* ~ 9*Hz*). We used the following parameter values: *C* = 1, *g_L_* = 0.25, *g*_1_ = 0.25, *τ*_1_ = 100. **c.** 2D linear system exhibiting damped oscillations in response to step-constant inputs (*f_nat_* ~ *f_res_* ~ 8*Hz*). We used the following parameter values: *C* = 1, *g_L_* = 0.05, *g*_1_ = 0.3, *τ*_1_ = 100.

**Figure 3:**
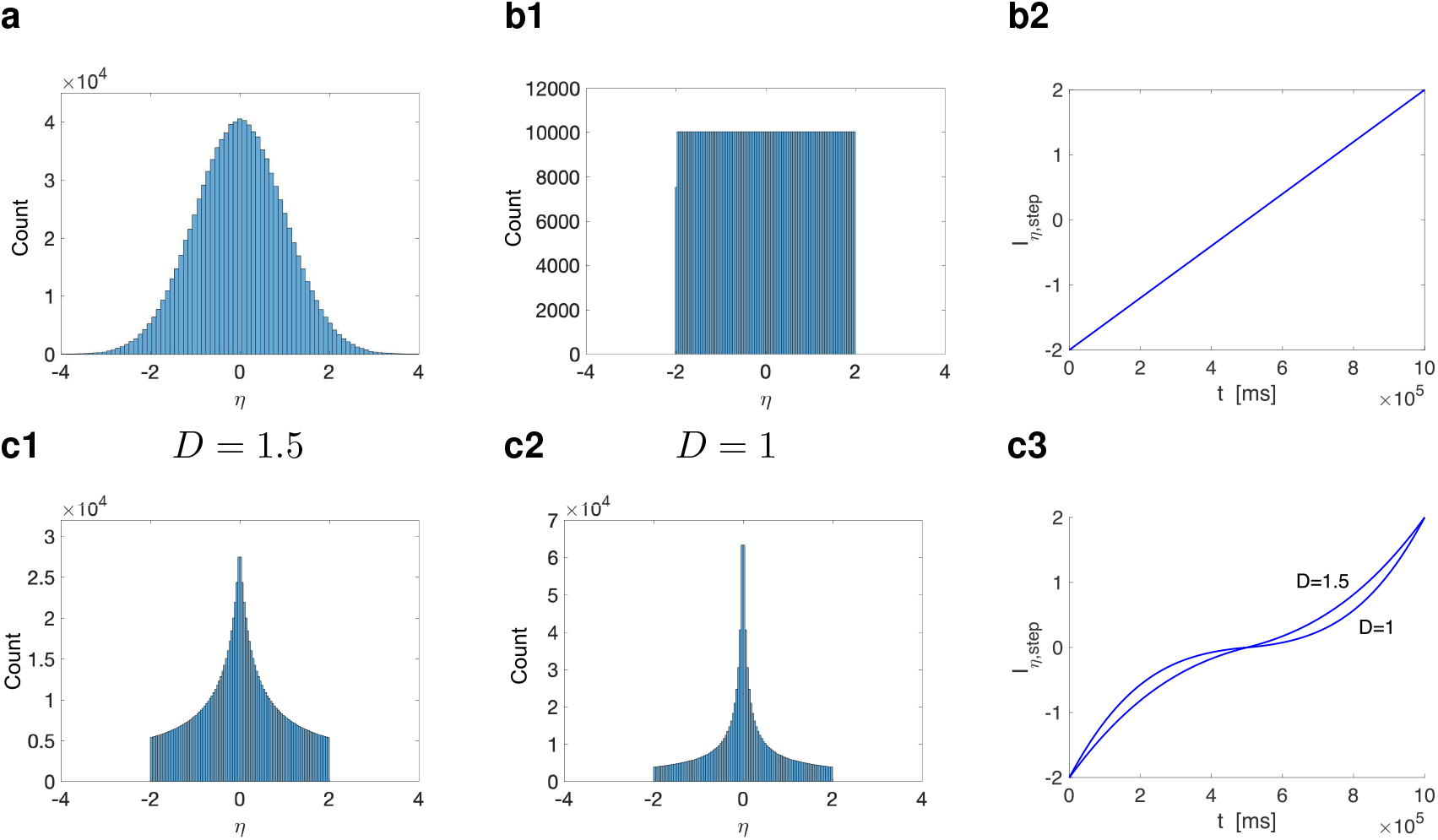
Histograms for representative distribution of the piecewise constant input amplitudes (*η*) of *I_η_*. To construct *I_η_*(*t*) we partition the time interval [0, *T_max_*] into *N* pieces of equal length Δ, and we define *I_η_*(*t*) = *η_k_* for *t* ∈ (*t*_*k* − 1_, *t_k_*) for *k* = 1, ... *N*. The values of the amplitudes 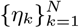 of the constant pieces are selected by following different rules (panels a to c). Each realization of *I_η_*(*t*) corresponds to a permutation of the order of the set 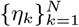. *I_η,step_*(*t*) is the special case where *η_k_* are organized in a non-decreasing order (used for reference). In all cases, Δ = 1 with a total time = 1000000 ms. **a.** Random normal distribution with mean zero and variance one. **b.** Equispaced (deterministic) distribution in the *η* interval [−2, 2]. *I_η,step_*(*t*) consist of the 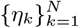 organized in an increasing order of amplitudes with c constant amplitude difference between adjacent pieces. **c.** Bell-shaped-like (deterministic) distribution in the *η* interval [*η_min_* = −2, *η_max_* = 2]. They were computed by (i) generating an equispaced distribution in the *η_aux_* interval [*η_min_, η_max_*], (ii) generating a cumulative distribution function (CDF) over *η_aux_* of a normal distribution with mean zero and variance *D*, (iii) computing *Amp_aux_* by flipping over (left to right) the left half the sequence of the CDF and multiplying it by 2 so to the resulting *Amp_aux_* lies in the interval [0, 1], (iv) Computing *Amp_int_* = *Amp_aux_* * (*η_max_* − *η_min_*)/(2 * Σ*Amp_aux_*). The right hand is computed by making it symmetric to the left hand over the ordinates axis. The ordered sequence 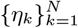 is computed from these amplitudes. **c1.***D* = 1.5. **c2.***D* = 1.

#### 2.2.2 Piecewise constant input functions with arbitrarily, but deterministically distributed amplitudes

The choice of normally distributed amplitudes (Fig. 3-a) is motivated by the fact that in the limit of Δ → 0, *I_η_* approaches white Gaussian noise when the piece duration goes to zero. In order to decouple the effect of the autonomous transient dynamics evoked by abrupt transitions between constant pieces and the randomness of the signal, we use fully deterministic distributions of amplitudes within some range. More specifically, we use arbitrary order permutations of a number of constant pieces initially arranged in increasing order of amplitudes where the amplitudes are chosen according to deterministic rules. Each protocol (see Section 2.2.1) consists of a subset of all possible permutations (of the order of constant pieces), which is the only source of uncertainty in the protocol.

We use two types of (deterministic) amplitude distributions for 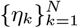 within the range [*η_min_, η_max_*]: equispaced (Fig. 3-b) and bell-shaped-like (Fig. 3-c). In the equispaced distribution the values of *η_k_* cover the full range [*η_min_, η_max_*] and satisfy *η*_*k* + 1_ – *η_k_* is equal for all *k* = 1, …, *N* − 1. The “ordered” PWC function *I_η,step_* is linear (Fig. 3-b2). The bell-shaped-like distribution was constructed from the random distribution (see the caption in Fig. 3). The corresponding “ordered” PWC function *I_η,step_* has an inflection point reflecting larger number of input amplitudes around zero (Fig. 3-c3).

### 2.3 Output Metrics

#### 2.3.1 Impedance (amplitude) profile

The impedance (amplitude) profile is defined as the magnitude of the ratio of the output (voltage) and input (current) Fourier transforms

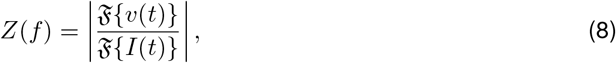

where 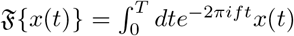. In practice, we use the Fast Fourier Transform algorithm (FFT) to compute 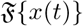. Note that *Z*(*f*) is typically used as the complex impedance, a quantity that has amplitude and phase. For simplicity, here we used the notation *Z*(*f*) for the impedance amplitude. Since we deal with discontinuous inputs (PWC), we have checked if Gibbs phenomenon would be an issue. As a result, we did not identify any ringing effect in the Fourier transforms related to Gibbs phenomenon.

#### 2.3.2 Voltage and impedance (amplitude) envelope profiles

The upper and lower envelope profiles 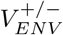 are curves joining the peaks and troughs of the steady state voltage response as a function of the input frequency *f*. The envelope impedance profile is defined as [17, 18]

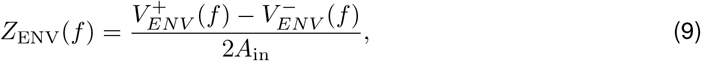

where *A*_in_ is the input amplitude. For linear systems, *Z*_ENV_(*f*) coincides with *Z*(*f*).

#### 2.3.3 Voltage power spectral density

In the frequency-domain, we compute the power spectral density (PSD) of the voltage as the absolute value of its Fourier transform 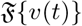. We will refer to this measure as PSD or *V*_PSD_.

#### 2.3.4 Peaks-and-troughs voltage response profiles

In each protocol, we compute the cells’ voltage response to the multiple trials described above. For each trial, we compute the sequence of the voltage response peaks and troughs consisting of the maximum value of the voltage *v* within the duration of a piece *η_k_* if *η_k_* > *η*_*k* − 1_ and the minimum of *v* within the range of a piece *η_k_* if *η_k_* < *η*_*k* − 1_, respectively. We use these peaks-and-troughs patterns or profiles (e.g., Fig. 4-a; difference between red and green dots in Fig 1 c-e) to characterize the voltage response patterns and their variability. We note that there are other possible metrics, including the whole *v* traces and the set of all peaks and troughs present in these traces (e.g., Fig. 4-c to -e). We found the peaks-and-troughs profiles as described above to be a simple and useful way to compare the voltage responses across trial (permutations).

**Figure 4:**
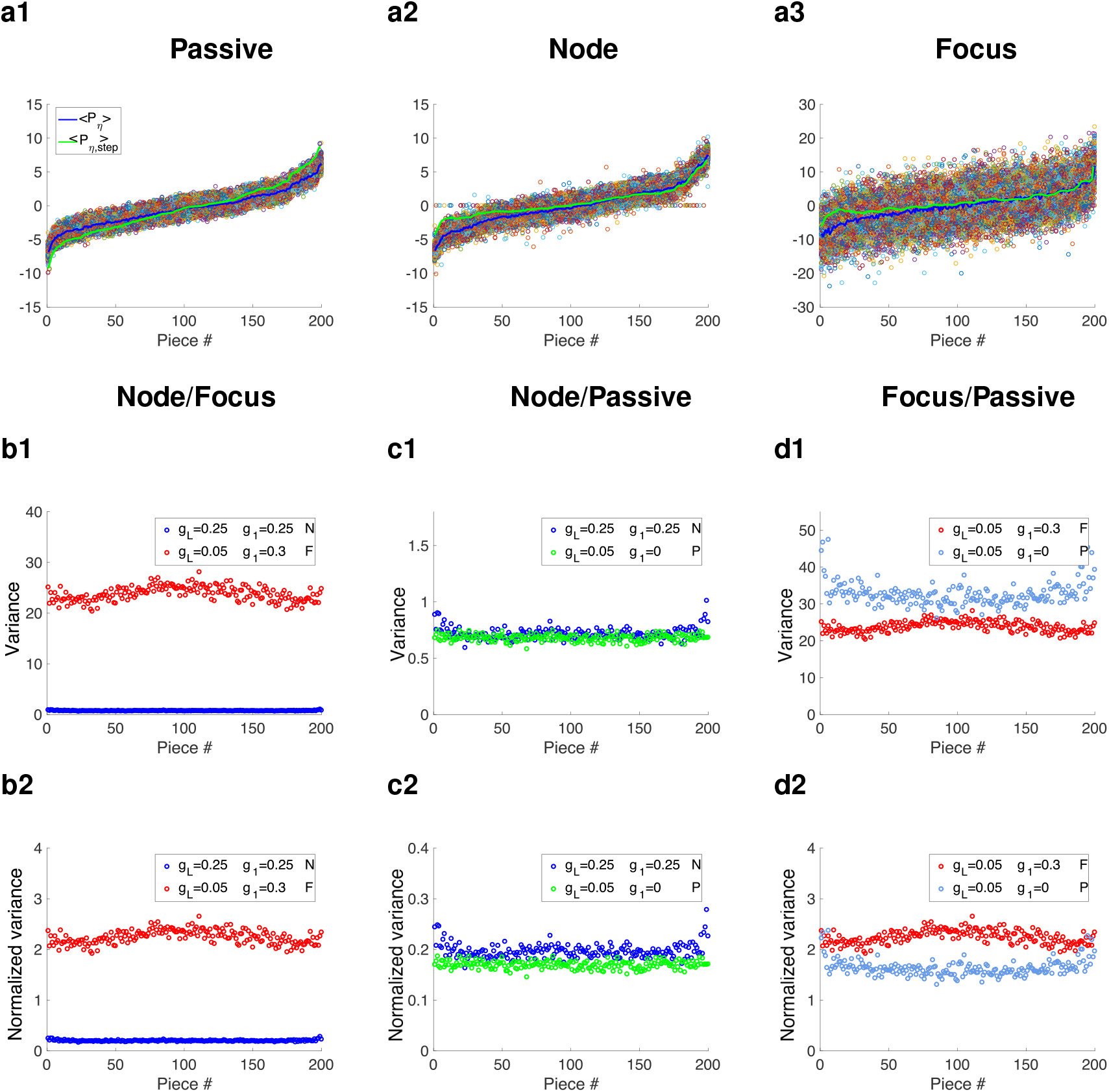
Variability (across trials) of the voltage responses of linear systems to piecewise constant inputs with normally distributed amplitudes. Trials consist of different permutations of the same set of constant pieces 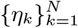. The piecewise constant inputs *I_η_* have Δ = 5, *N* = 200 (a, b) and *N* = 2000 **(a)**. The parameter values are the same as in Fig. 1. The color dots indicate the peaks-and-troughs patterns for all trials reorganized so that the corresponding values of *η_k_* from which they originate are ordered in a monotonically increasing manner. All dots for a given piece correspond to the same value of *η_k_*. *< P_η_ >* (blue) is the mean value of the reordered peaks-and-troughs patterns for each linear piece. *< P_η,step_ >* (green) is the peaks-and-troughs pattern corresponding to the (ordered) input function *I_η,step_*. **b1 to d1.** Variance (b1 to d1) of the peaks-and-troughs patterns in panel a. **b1.** Comparison between cells having a node and a focus. **c1.** Comparison between a cell having a node and the corresponding passive cell. **d1.** Comparison between a cell having a focus and the corresponding passive cell. **b2 to d2.** Normalized variance (b2 to d2) computed as the variance (b1 to d1) divided by the peak of the unforced cells’ response to a step-constant input of amplitude 1. **b2.** Comparison between cells having a node and a focus. **c2.** Comparison between a cell having a node and the corresponding passive cell. **d2.** Comparison between a cell having a focus and the corresponding passive cell. We used the following parameter values: (i) *C* = 1, *g_L_* = 0.25, *g*_1_ = 0.25, *τ*_1_ = 100 for the node, and (ii) *C* = 1, *g_L_* = 0.05, *g*_1_ = 0.3, *τ*_1_ = 100 for the focus.

#### 2.3.5 Rearranged peaks-and-troughs voltage response profiles

In order to compare the voltage responses across trials we rearrange the peak-and-troughs voltage response profiles according to the increasing amplitude values of the elements of the set 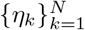 from which they were evoked. We use the notation *P_η_* for the resulting rearranged voltage response patterns and *P_η,step_* for these voltage response profiles produced by *I_η,step_* (inputs with increasing order of constant piece amplitudes). In this way, we compare multiple different ways in which the response to a given constant piece is produced by transitions from pieces with different amplitudes (i.e., different initial conditions with respect to that constant piece). The differences in the voltage peak/troughs values across trials for each piece will be due to the differences in the values of *V* and the other variables at the arrival time of that piece (initial conditions) and therefore to the differences in the transient dynamics that they evoke.

#### 2.3.6 Numerical simulations

We used the modified Euler method (Runge-Kutta, order 2) [81] with step size Δ*t* = 0.01 ms. Preliminary simulations where lower step sizes than this have been tested did not change our results. All neural models and metrics, including phase-plane analysis, were implemented by self-developed Matlab routines (The Mathworks, Natick, MA) and are available in https://github.com/BioDatanamics-Lab/impedance_input_dependent.

## 3 Results

### 3.1 Voltage response to piecewise constant inputs with variable amplitudes: sequence of autonomous transient dynamics followed by steady-state responses

Figs. 1-c1 to -e1 illustrate the voltage response of three types of cells to the same PWC inputs with increasing amplitude (generated by *I_step_*, see Fig. 1-a, bottom). We use long piece durations here only for explanatory purposes. The voltage response to each linear piece consists of a transient and a steady-state components, which depend on the model parameters. The properties of the transient component in each case (the autonomous transient dynamics) are qualitatively different for the three cell types. Passive cells (and cells behaving as passive cells) (Fig. 1-c1) exhibit a monotonic increase towards the steady-state. The node cells (cells having a stable node, N-cells) in Fig. 1-d1 display overshoots and the focus cells (cells having a stable focus, F-cells) in Fig. 1-e1 display damped oscillations before converging to the steady-state. We note that not all two-dimensional cells having a stable node or focus show prominent overshoots or damped oscillations.

Figs. 1-c2 to -e2 show the voltage response of the same three cell types to PWC inputs with the same constant pieces as in Figs. 1-c1 to -e1, but the three intermediate pieces are arranged in a different order. The type of autonomous transient dynamics, which are activated at the transition points between input pieces, remains the same as in Figs. 1-c1 to -e1, but the response amplitude of the voltage response pattern increases for the N- and F-cells as compared to Figs. 1-d1 and -e1. This is due to a combination of factors that involve the differences in the jump sizes between linear pieces and the differences in initial conditions for each linear piece (e.g., Fig. 1-b). The values of the participating variables prior to the transition between linear pieces serve as initial conditions for the new regime. While for 1D system, the response amplitude is bounded by the steady-state for each linear piece regardless of the initial conditions, for 2D systems the amplitude response depends on the initial conditions not only of the main variable (*v*), but also the recovery variable that is not shown in the graph (e.g., Fig. 1-b).

### 3.2 Oscillatory voltage response properties to piecewise constant inputs with arbitrarily distributed amplitudes

The *V* patterns discussed above (Fig. 1, row 2) for a small number of input (*I_η_*) partitions (large Δ = 200) display the fully developed autonomous transient dynamics of the three cell types to step-constant inputs. The extension of these results to the *V* patterns generated by inputs with a larger, more realistic number of partitions (smaller Δ) is not straightforward since the smaller the partition size the less time the cell has to develop the autonomous transient behavior corresponding to each partition. Still, the transient dynamics reflect the model properties and the differences among models.

#### 3.2.1 Piecewise constant inputs with normally distributed amplitudes and a large enough number of partitions uncovers the oscillatory dynamic properties of the target cells

Figs. 2 shows the *V* response patterns to inputs *I_η_* with a larger number of partitions (Δ = 5). Figs. 2-a1 and -b1 show no apparent oscillatory pattern, while Fig. 2-c1 shows irregular oscillations, which is confirmed by the PSD graph in Fig. 2-c2. The PSD graph in Fig. 2-b2 uncovers the presence of oscillations in the corresponding *V* pattern (Fig. 2-b1), while the PSD graph in Fig. 2-a2 confirms the absence of oscillations consistent with the 1D dynamics of the cell. Interestingly, not only the F-cell that shows damped voltage oscillations in response to step-constant inputs shows oscillatory activity in response to *I_η_*, but also the N-cell that shows overshoots in response to step-constant inputs. This would be in principle not surprising since both cell types show resonance in response to sinusoidal inputs [16, 17], but the PSDs for the input patterns *I_η_* with a smaller number of partitions (Δ = 200 as in Fig. 1, row 3) shows no oscillations for the three responses (see insets in Fig. 2, row 2). The arbitrarily (randomly) distributed amplitudes allow the voltage response to explore a wide region of the phase-space and therefore capture information about the vector field governing the transient dynamics. The short duration of the pulses keeps the trajectory (“almost continuously") exploring the vector field by sequentially activating the autonomous transient dynamics, and this “permanent motion” prevents the interference of the regions of the vector field governing the dynamics in close vicinities of the steady-state. This mechanism is conceptually similar to the one governing the generation of resonance in response to oscillatory inputs in both linear and nonlinear systems [17, 18] where resonance can be observed in the absence of intrinsic oscillatory behavior [16, 17, 21].

#### 3.2.2 Uncovering the oscillatory dynamic properties of the target cells requires constant pieces with arbitrarily distributed amplitudes, but not amplitude randomness

Here we decouple the effects of the autonomous transient dynamics (in response to the sequence of input constant pieces) from the pieces’ amplitude randomness on the ability of these PWC inputs to uncover the oscillatory (intrinsic) and resonant properties of the receiving cells.

To this end we use fully deterministic distributions of amplitudes within some range as described in Section 2.2.2. This leaves the choice of the subset of all possible permutations (number of trials) for each protocol as the only source of uncertainty in the input signal. Fig. S1 (Δ = 5 and Δ = 1) show the response patterns to these PWC inputs (equispaced distributed, deterministic amplitudes) for the same parameter values as Fig. 2 (random amplitudes). The *Z*-profiles in these three figures are almost identical. The *V* PSD profiles in Figs. 2 and S1 are also very similar. The differences between the *V* PSD profiles in these figures and the ones for Δ = 5 are due to the larger value of the total time used there. Together, these results show that randomness is not needed to uncover the resonant and oscillatory properties of the receiving cells and these oscillatory properties emerge almost exclusively as the result of the sequential and fast activation of the cell’s autonomous transient dynamics.

#### 3.2.3 PWC inputs with arbitrarily distributed amplitudes capture the nonlinear amplification of the oscillatory voltage responses

The amplification of the voltage response oscillations to PWC inputs discussed so far was presented in the context of linear systems as the result of changes in the model parameter values (e.g, from N- to F-cells in Figs. 2-, S1-, -b and -c). Because the models used there are linear, the dependence of the voltage response amplitude on the input amplitude (*D*) is simple. Increasing values of *D* cause proportional increases in the voltage response amplitude so that the *Z*-profile remains unchanged.

Here we focus on the nonlinear amplifications produced in (nonlinear) models as the result of increasing values of the input amplitude *D*. To this end we use the piecewise linear (PWL) model (3)-(5), which is a continuous, nonlinear extension of the linear model (1)-(2) where the *V* -nullcline is “broken” (e.g., Figs. 5-b). It was shown in [17] that this type of models display nonlinear amplifications of the voltage response to sinusoidal inputs and capture similar phenomena observed in more realistic nonlinear models, in particular two-dimensional models having parabolic-like *V* -nullclines describing the subthreshold voltage dynamics. [18].

**Figure 5:**
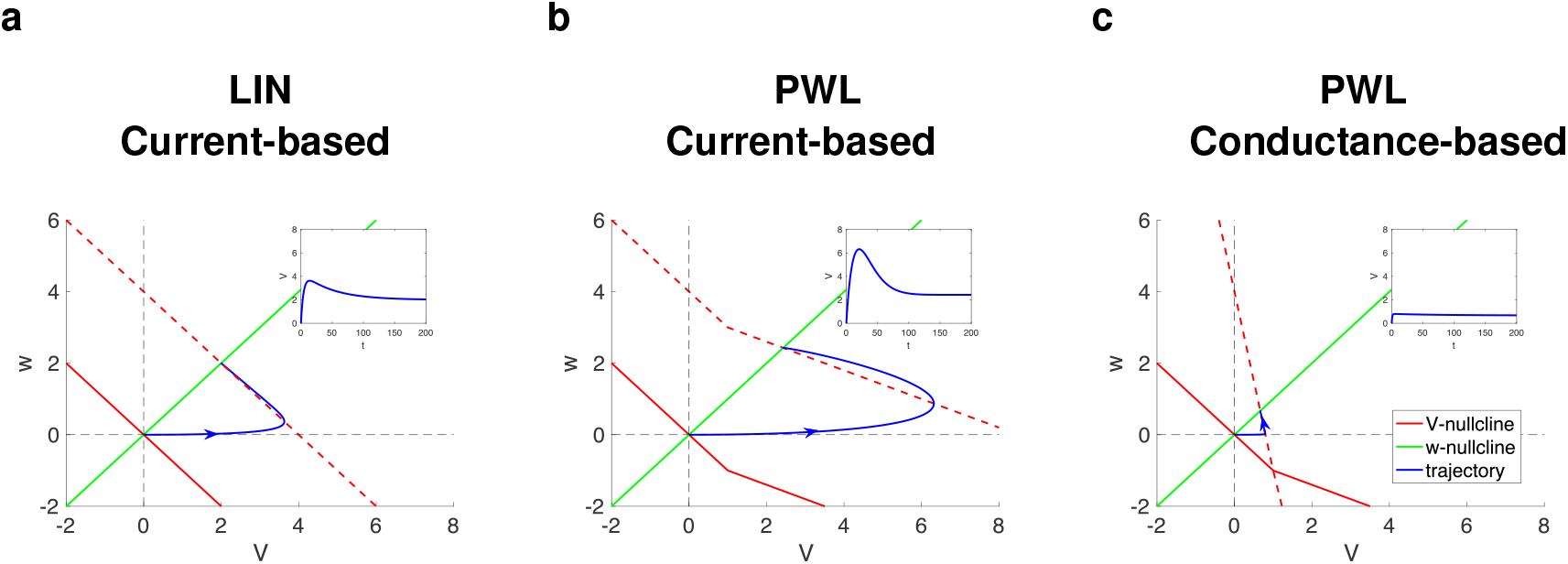
Nonlinear transient voltage response amplifications and attenuations in current-vs. conductance-based inputs. Phase-plane diagrams for *I* = 1. The solid-red curve represents the *V* -nullcline for *I* = 0, The dashed-red curve represents the *V* -nullcline for *I* = 1, the solid-green curve represents the *w*-nullcline for *I* = 0, and the solid-blue curve represents the trajectory which initially starts at (0, 0) (the fixed-point for *I* = 0) and converges to the fixed-point for *I* = 1. The insets show the *V* traces. The 2D linear system exhibits and overshoot in response to step-constant inputs and resonance in response to oscillatory inputs [16, 17, 21]. **a.** Linear (LIN) model described by eqs. (1)-(2). **b.** Current-based piecewise linear (PWL) model described by eqs. (3)-(5). **c.** Conductance-based piecewise linear (PWL) model described by eqs. (6)-(7) and (5) with *S* substituted by *I*. We used the following parameter values: *C* = 1, *g_L_* = 0.25, *g*_1_ = 0.25, *τ*_1_ = 100 (same as in Figs. 1 and 2-b), *v_c_* = 1 and *g_c_* = 0.1.

We first introduce the ideas by examining the autonomous transient dynamics of the PWL model in response to a constant input *I* and then use these results to understand the nonlinear amplification of this PWL model in response to PWC inputs with increasing values of *D*. Figs. 5-a and -b show the superimposed phase-plane diagrams for a PWL model (b) and the linear model (a) from which it originates for *I* = 0 (solid-red, baseline) and *I* = 1 (dashed-red). The *w*-nullclinne is unaffected by changes in *I*. The trajectory (blue), initially at the fixed-point for *I* = 0, evolves towards the fixed-point for *I* = 1. In both cases (Figs. 5-a and -b) the voltage response exhibits an overshoot. The peaks (inset) occur when the trajectories cross the *V* -nullcline. For low enough values of *I* (lower than in Figs. 5-a and -b) the trajectory remains within the linear region (the trajectory does not reach the *V* -nullcline’s “breaking point” value of *V*) and therefore the increasing values of *I* produce a linear voltage response amplification (no differences between the responses of the linear and PWL model; not shown). The nonlinear amplification is apparent for values of *I* for which the *v*-nullcline is high enough so the trajectory is able to cross from one linear regime (determined by the left piece of the dashed-red V-nullcline) where the trajectory is at t=0, to the other (determined by the right piece of the dashed-red V-nullcline). The virtual fixed-point moved from the position where *I* = 0 and converged to the position where nullclines cross for *I* = 1. Because the *V* -nullcline’s “right piece” has a smaller slope than the “left piece” (the slope it would have if it were linear), the trajectory is able to reach larger values of *V* for the nonlinear system than for the corresponding linear one before turning around, and therefore the peak for the nonlinear system is higher than for the linear system. Nonlinear voltage response amplifications in this type of system are dependent on the time scale separation between the participating variables. For smaller values of *τ*_1_ this nonlinear amplification is reduced and although the system is nonlinear, it behaves quasi-linearly [17, 18].

The nonlinear amplification discussed above is particularly stronger for the transient dynamics (initial upstroke) than for the steady-state response, and therefore it is expected to have consequences for the nonlinear responses of nonlinear models to PWC inputs with large enough values of *D* (Fig. 6). We use a PWC input with a (non-random) equispaced distribution of constant piece amplitudes in the range [0, 2] multiplied by *D*. For small enough values of *D* (Fig. 6-a1) the trajectory remains within the linear regime and therefore the responses to the linear and the PWL models are almost identical (blue and red). As *D* increases, the nonlinear amplification becomes stronger (Figs. 6-b1 and -c1, blue and red). This is accompanied by similar changes in the *Z* profile (not shown). As discussed above, these mechanisms are dependent on the time scale separation between the participating variables, determined by *τ*_1_ and the nonlinear voltage response amplification is reduced for smaller values of *τ*_1_ (see Fig 6-a2, -b2, and - c2 for *τ*_1_=10). Lowering the parameter *τ*_1_ makes the system more linear and reduces the amplification.

**Figure 6:**
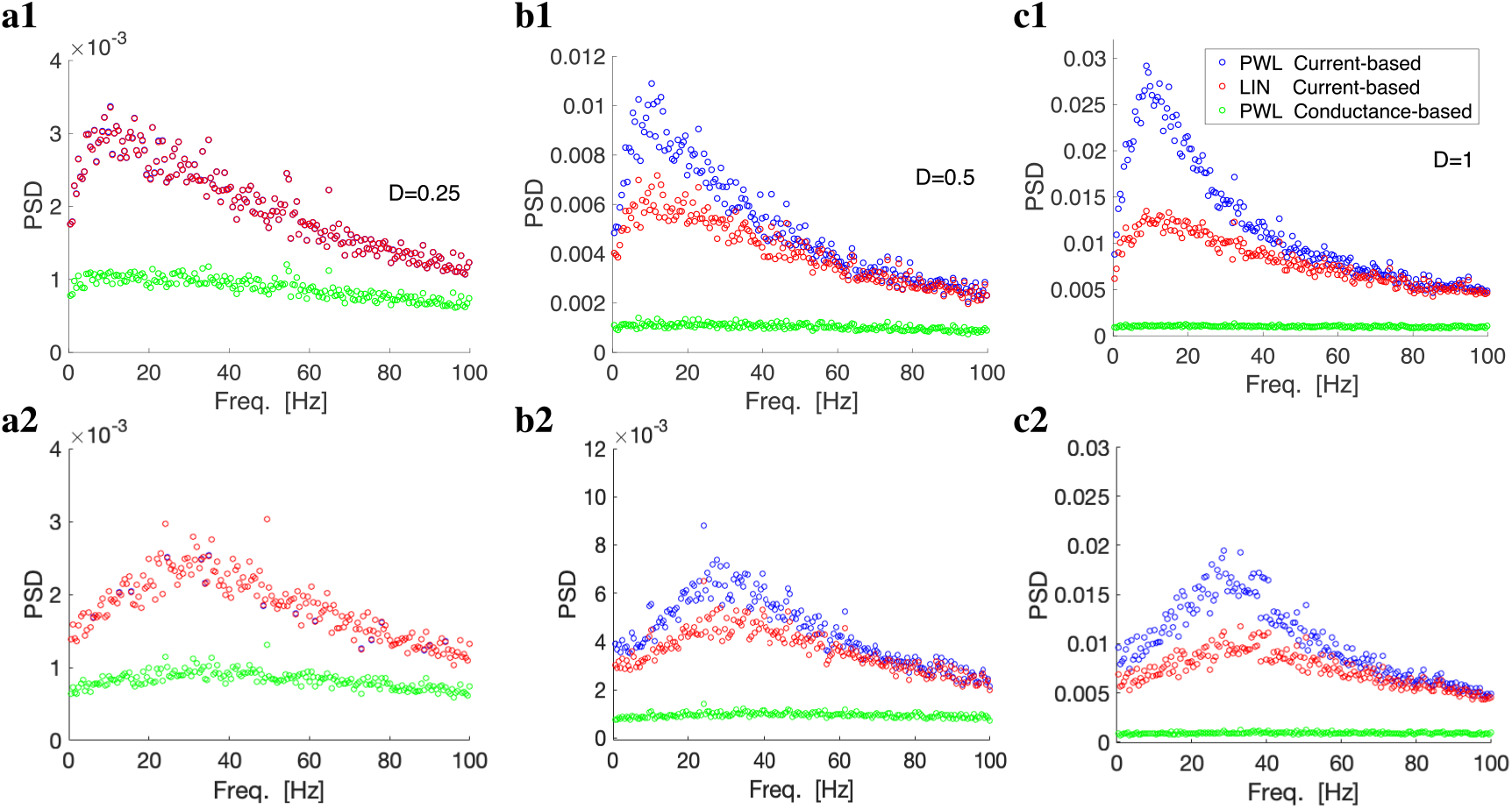
Oscillatory voltage responses for current-vs. conductance-based piecewise constant inputs with equispaced (non-randomly) distributed amplitudes and different time constants. The piecewise constant inputs *I_η_* have Δ = 1 (total time = 1000000 ms, N = 1000000). The constant pieces’ amplitudes are equispaced and deterministically distributed in the range [0, 2]. The 2D linear system exhibits and overshoot in response to step-constant inputs and resonance in response to oscillatory inputs. Parameter values are as in the first row of Fig. 6 (the parameter values for the linear systems are the same as in Figs. 1 and 2-b). Power spectra density (PSD) profiles for a sample *V* trace. The piecewise constant inputs *I_η_* (total time = 1000000 ms, N = 1000000) have different values of *D*. **a.** *D* = 0.25. **b.** *D* = 0.5. **c.** *D* = 1. We used the following parameter values in **a1**, **b1.**, and **c1.**: *C* = 1, *g_L_* = 0.25, *g*_1_ = 0.25, *τ*_1_ = 100, *v_c_* = 1 and *g_c_* = 0.1. We used the following parameter values in **a2**, **b2.**, and **c2.**: *C* = 1, *g_L_* = 0.25, *g*_1_ = 0.25, *τ*_1_ = 10, *v_c_* = 1 and *g_c_* = 0.1.

#### 3.2.4 Oscillation amplification and attenuation: current- versus conductance-based responses to synaptic-like PWC inputs

The oscillatory dynamics considered above emerge in response to additive PWC current inputs. However, the synaptic inputs received by neurons are multiplicative and conductance-based as described by the model (6)-(7).

From the phase-plane diagram in Fig. 5-c we see that increasing values of *I* (replacing *S* in the model) reduces the nonlinearity of the *V* -nullcline (dashed-red) and increases (in absolute value) its slope. Both phenomena oppose the voltage response amplification (blue) and the overshoot becomes much less prominent. The triangular region (bounded by the *V* -axis, the displaced *V* - nullcline (dashed-red) and the *w*-nullcline (green)) is reduced in size as compared to the current-based inputs (panel b) and therefore the response is reduced in amplitude. Moreover, because the displaced *V* -nullcline in panel c is more vertical than the baseline *V* -nullcline, the size of the voltage overshoot in response to constant inputs is reduced and, in this sense, the voltage responses become quasi-1D. As a consequence, the initial portion of the transient responses to abrupt changes in input is reduced in size. This translates into the oscillatory response to PWC inputs, which is also attenuated and the resonant peak disappears or is significantly reduced (Figs. 6-c1, green).

### 3.3 Emergence of variability in response to piecewise constant inputs with normally distributed amplitudes

Here we address the relationship between the transient dynamic properties of individual cells (autonomous transient dynamics) and the variability of their responses to piecewise constant (PWC) input functions with pieces of the same duration and randomly distributed amplitudes. We primarily use Gaussian distributions and a relatively large number of pieces (with a small duration each). This allows us to separate the effects of the autonomous transient dynamics activated by the transition between pieces with variable size-steps from the variability of the piece duration, which is chosen to be in a range much lower than the cell’s intrinsic (natural) and resonant frequencies. In the next Section, we show that amplitude randomness is not needed, but only the arbitrary order of the amplitude of the constant pieces.

#### 3.3.1 Variability in response to piecewise constant inputs emerges from the transient response properties of the autonomous system to step constant inputs

The voltage response of cells to PWC inputs consists of a sequence of transient behaviors (initiated immediately after the transition between two constant pieces), reflecting the autonomous transient dynamics in response to a set of initial conditions for the participating variables, followed by an approximation to the steady-state for each input piece. The latter and part of the former may be absent if the duration Δ of each constant piece is not long enough. As discussed above, Figs. 1-c1 to -e1 illustrate three qualitatively different ways in which cells respond to a sequence of increasing step function inputs according to whether they have a stable node (N, 2D), a stable focus (F, 2D) or they are passive (P, 1D). For illustrative purposes, the value of Δ was chosen to be large enough so as to show both dynamic components of the response to each linear piece. In subsequent simulations, Δ will be chosen to be much smaller so as to capture realistic situations (as discussed above), and therefore the voltage response will capture only the initial portions of the autonomous transient dynamics.

Passive (1D) cells exhibit a monotonic behavior towards the steady-state in response to each input piece. For the parameters chosen, the transient increase is relatively fast (Fig. 1-c1). 2D cells, in addition, can display overshoots (Fig. 1-d1, N) and damped oscillations (Fig. 1-e1, F) as they approach the steady-state in response to each input piece. The peak amplitudes of *V* in response to each input piece correspond to the transient component of the response and depend on the initial conditions in that regime (Fig. 1-b), which in turn depend on the values both *V* and particularly *w* reach at the end of the previous regime. Because of this sensitivity of the transient responses to initial conditions, an order rearrangement of the constant pieces produces different response patterns (Figs. 1-c2 to -e2). This variability, which is reflected in the peak and trough values of the “disordered” patterns (c2, d2, e2) as compared to the “ordered” ones (c1, d1, e1) is due to the differences in the initial conditions in each regime as *v* transitions between pieces with different amplitudes. The variability among patterns generated by different permutations of the set 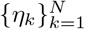 is inherited from this principle since the initial conditions for a given input piece depends on the “last” values of the participating variables in response to the previous piece. The variability is stronger for the 2D cells (Fig. 1-d and Fig. 1-e) than to for the 1D cell (Fig. 1-c1) since the sensitivity to initial conditions is weaker for the former than for the latter. The addition of a second variable (dimension) typically adds sensitivity to initial conditions.

#### 3.3.2 The response variability to piecewise constant inputs with normally distributed amplitudes depends on the cell’s autonomous transient dynamic complexity

Fig. 4-a (colored dots) shows the distribution of the peaks-and-troughs patterns *P_η_* for 100 trials (permutations of a randomly distributed set 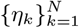, *N* = 200, Δ = 5) for a passive cell (P), a cell having a node (N) and a cell having a focus (F). The average *< P_η_ >* and the *P_η,step_* curves, superimposed to *P_η_*, show that the average across trials and the response to the reference pattern (with minimal variability) are not identical. In this figure they are relatively close.

The variability of *P_η_* across trials, Var(*P_η_*) is larger for the F-cell than for the N-cell and there seems to be no significant difference between the variability of N- and P-cells. (The latter two are comparable since they have the same value of *g_L_*.) However, the qualitative differences between N- and F-cells are accompanied by different strengths in the responses to step-constant inputs (e.g., Fig. 1), and one has to account for these differences in order to isolate the effects of the type of transient dynamics. We note that these parameter values were chosen so that the two cells (N and F) have resonance in the same frequency band.

Figs. 4-b1 to -d1 show the variances of *P_η_* (Var(*P_η_*)) and Figs. 4-b2 to -d2 show the normalized variances (VarN(*P_η_*)) computed as these variances divided by the peak of the unforced cells’ response to a step-constant input of amplitude 1. Fig. 4-b show that both Var(*P_η_*) and VarN(*P_η_*) are larger for the F-cell than for the N-cell. Fig. 4-c shows that both Var(*P_η_*) and VarN(*P_η_*) are slightly larger for the N-cell than for the corresponding P-cell (obtained by making *g*_1_ = 0). In contrast to this, Fig. 4-d shows that Var(*P_η_*) and VarN(*P_η_*) have different relative values for the F-cell and the corresponding P-cell; Var(*P_η_*) is larger for the P-than for the F-cell, but VarN(*P_η_*) is larger for the F-than for the P-cell. The values of these quantities in both cases are significantly larger for the F-/P-cells than for the N-/P-cells. Together, these results suggest that under the constraints imposed by the two cells having resonance in the same frequency band, the voltage response of the F-cell is more variable than the voltage response of the N-cell, and the larger variability of the F-/P-cells as compared to the N-/P-cells is due to a smaller value of *g_L_*, which in turn indicates a stronger amplification of the transient voltage response to step-constant inputs. However, the larger variabilities cannot be attributed to these differences in the voltage response amplitudes since they persist after the variances have been normalized.

A similar result is obtained when one lifts the resonance constraint on the N- and F-cells. Fig. 7-b shows the effects of changes in *g*_1_ on Var(*P_η_*) and VarN(*P_η_*) for fixed values of *g_L_*. As *g*_1_ increases the transient dynamics of the cell transitions from P (green) to N (blue, *f_res_* = 7) to F (red, *f_res_* = 14, *f_nat_* = 12.3) (Fig. 7-a). Var(*P_η_*) is slightly larger for the F-than for the N-cell and larger than these two for the P-cell. In contrast, VarN(*P_η_*) is much larger for the F-than for the N-cell, and VarN(*P_η_*) for the P-cell is comparable to that for the N-cell.

**Figure 7:**
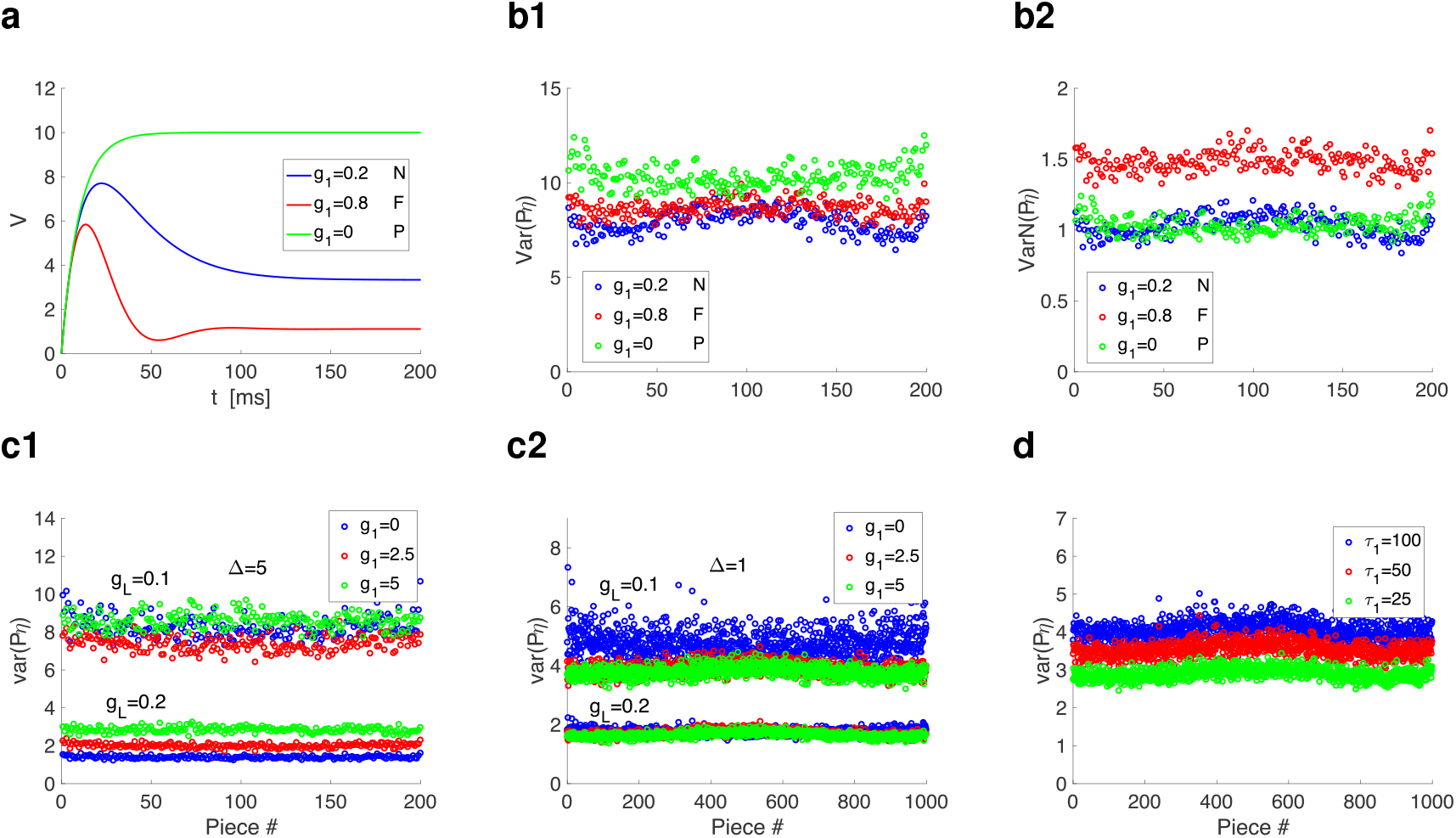
Variability (across trials) of the responses of linear systems to piecewise constant inputs with normally distributed amplitudes. Trials consist of different permutations of the same set of constant pieces 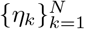. The piecewise constant inputs *I_η_* have Δ = 5, *N* = 200. **a.** *V* traces for the node, focus and passive cells in response to a constant input of amplitude 1. **b.** Effects of changes of the linearized resonant conductance *g*_1_ on the variances Var(*P_η_*) and normalized variances VarN(*P_η_*) of the peaks-and-troughs patterns. Normalized variance computed as the variance (b1) divided by the peak of the unforced cells’ response to a step-constant input of amplitude 1 (a). **b1.** Var(*P_η_*). **b2.** VarN(*P_η_*). We used the following parameter values: *C* = 1, *g_L_* = 0.1, *τ*_1_ = 100, *g*_1_ = 0.2 (node), *g*_1_ = 0.8 (focus) and *g*_1_ = 0 (passive). **c.** Effects of changes of the linearized conductances *g_L_* and *g*_1_ on the variances Var(*P_η_*). **c1.** Δ = 5. **c2.**Δ = 1. We used the following additional parameter values: *C* = 1 and *τ*_1_ = 100. **d.** Effects of changes of *τ*_1_ on the variances Var(*P_η_*). We used Δ = 1. We used the following additional parameter values: *C* = 1, *g_L_* = 0.1 and *g*_1_ = 2.

#### 3.3.3 The response variability along time and across trials depends on the levels of complexity of the autonomous transient dynamics

If the voltage response variability to PWC inputs with randomly distributed amplitudes primarily depends on the receiving cell’s autonomous transient dynamics, then one expects the variability to be higher, the faster the response of the individual cells to step-constant inputs. As this response becomes faster, then it is easier for *V* to reach values further away from the mean The parameter *g_L_* is the ideal one to test these ideas since it determines the time constant of the *V* equation. The smaller *G_L_*, the faster the response. Fig. 7-c shows that Var(*P_η_*) is higher for *g_L_* = 0.1 than for *g_L_* = 0.2 and this remains true for various representative values of *g*_1_. The dependence of Var(*P_η_*) with *g*_1_ is more complex and less clear. For *g_L_* = 0.2, Var(*P_η_*) increases with *g*_1_, while for *g_L_* = 0.1, Var(*P_η_*) first decreases and then increases with increasing values of *g*_1_. For Δ = 1 these dependences are less well separated (Fig. 7-c2) when compared with Δ = 5 (Fig 7-c1).

For fixed values of *g_L_* and *g*_1_, the time constant *τ*_1_ associated to the recovery variable *w* controls the time separation between the variables *v* and *w*. The larger *τ*_1_ (the slower *w*), the faster the autonomous transient response. Because of this stronger sensitivity to initial conditions, the variability is expected to be larger, which is confirmed by Fig. 7-d.

Together, these results show how the variability of the responses to PWC input functions with normally distributed amplitudes are controlled by the transient dynamics of the autonomous responses to piecewise constant inputs. This variability emerges as the input pieces within the same set are permuted for different trials. The differences among the responses to the same piece (piece with the same amplitude) across trials are due to differences in the initial conditions relative to these pieces caused primarily by the varying amplitudes of the preceding pieces.

#### 3.3.4 The peak-and-trough profiles are able to capture the nonlinear properties of the target cells

In our previous discussion, we have used linear models as the receiving cells to the PWC inputs. We have also shown that the presence of certain types of nonlinearities affects not only the vector field but also the effective time constants of the cell, which in turn affect their autonomous transient dynamics. We reasoned that these types of nonlinearities may also affect the variability of the cells’ response to PWC inputs. To test these ideas we used the piecewise-linear (PWL) model (3)-(5) where the *v*-nullcline (right hand side of the first equation equal to zero) is broken into two linear pieces at *v* = *v_c_*. We chose *v_c_* > 0 to be within the range of values of the *v* response to the PWC input and *g_L_ > g_c_*. Effectively, the cell’s membrane time constant transitions from 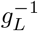 to 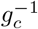 at *v* = *v_c_*. Therefore one should expect the variability pattern to be larger. Since the PWL model is asymmetric with respect to the equilibrium of the autonomous cell, then one should expect the variability pattern to be asymmetric with respect to the responses mean. Figs. 8 (rows 1 and 2) illustrate these ideas. Figs. 8 (row 3) summarize these results and also shows that the variability for the nonlinear models shows a strong dependence on the amplitude of the input constant pieces, while the variability pattern for the linear model is flatter. This is also expected since the larger the input amplitude, the more likely the response to reach values beyond *v_c_*. However, the presence of nonlinearities in the model does not necessarily guarantee a nonlinear response to external inputs. The latter strongly depends on the time scale separation between the participating variables as it occurs in other types of responses (e.g., to oscillatory inputs) where the mechanisms depend on the cell’s autonomous transient dynamics [18]. The stronger the time scale separation (the larger *τ*_1_), the stronger the nonlinear amplification of the voltage response to time-dependent inputs. Consistent with this, Fig. 8-c1 shows that the peaks-and-troughs profiles for the nonlinear system and *τ*_1_ = 10 is more symmetric and spans a smaller range than the profile for *τ*_1_ = 100 (Fig. 8-b1), while the peaks-and-troughs profiles for the corresponding linear systems are similar (Fig. 8-b2 and -c2). In addition, Var(*P_η_*) is smaller and flatter for *τ*_1_ = 10 (Fig. 8-c1) than for *τ*_1_ = 100 (Fig. 8-c2).

**Figure 8:**
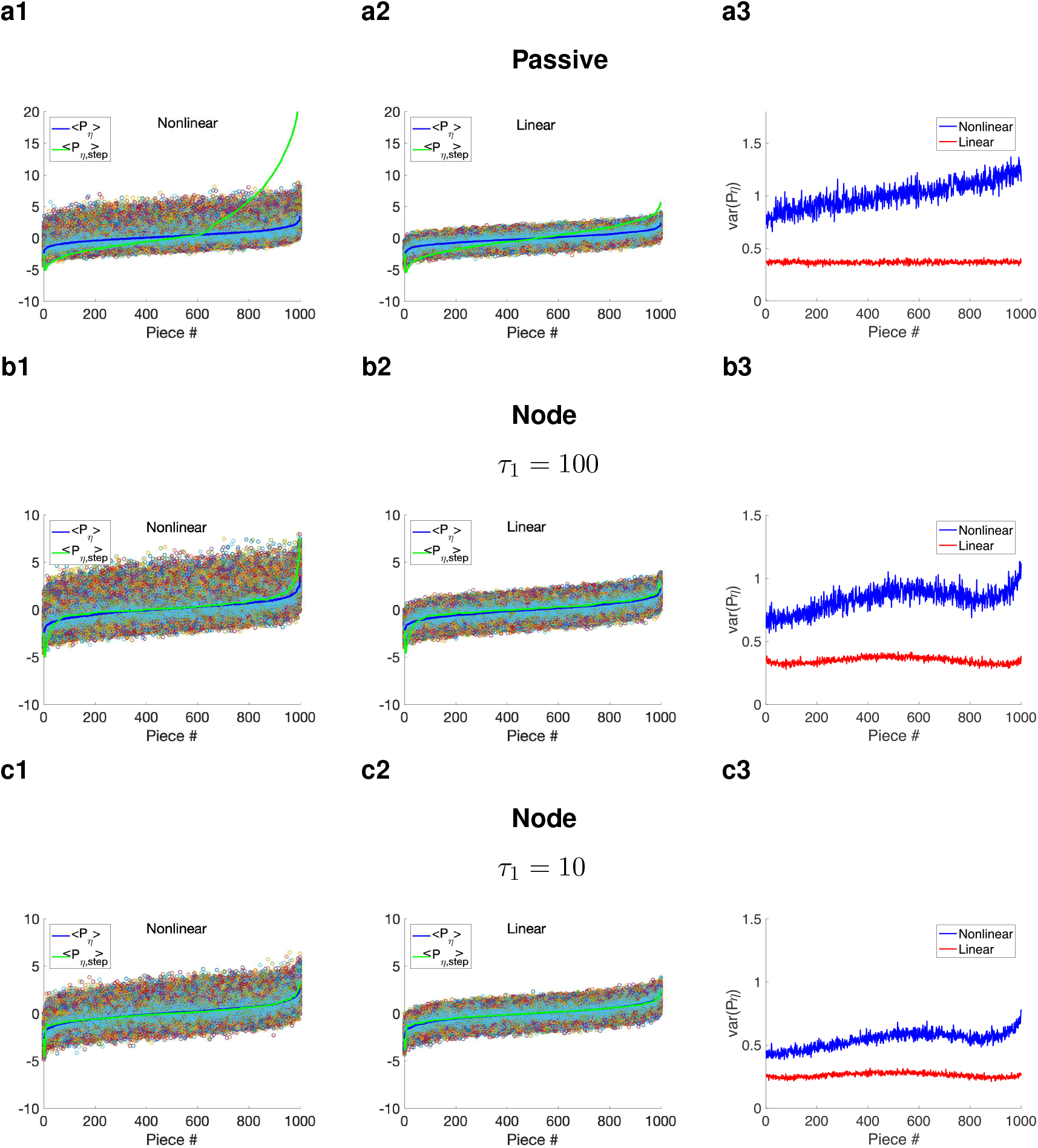
Effects of the model nonlinearities on the variability (across trials) in response to linear systems to piecewise constant inputs with normally distributed amplitudes. Trials consist of different permutations (1000) of the same set of constant pieces 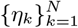. The piecewise constant inputs *I_η_* have Δ = 1, *N* = 1000. The piecewise linear model is given by (3)-(4). **Left and middle.** The color dots indicate the peaks-and-troughs patterns for all trials reorganized so that the corresponding values of *η_k_* from which they originate are ordered in a monotonically increasing manner. All dots for a given piece correspond to the same value of *η_k_*. *< P_η_ >* (blue) is the mean value of the reordered peaks-and-troughs patterns for each linear piece. *< P_η,step_ >* (green) is the peaks-and-troughs pattern corresponding to the (ordered) input function *I_η,step_*. **Right.** Comparison of Var(*P_η_*) between the linear (red) and piecewise linear model (blue). **a.** We used the following parameter values: *C* = 1, *g_L_* = 0.5, *g*_1_ = 0, *g_c_* = 0.1 and *v_c_* = 0.5. **b.** We used the following parameter values: *C* = 1, *g_L_* = 0.5, *τ*_1_ = 100, *g*_1_ = 1, *g_c_* = 0.1 and *v_c_* = 0.5. **c.** We used the following parameter values: *C* = 1, *g_L_* = 0.5, *τ*_1_ = 10, *g*_1_ = 1, *g_c_* = 0.1 and *v_c_* = 0.5.

### 3.4 The variability in the response to piecewise constant inputs requires constant pieces with arbitrarily distributed amplitudes, but not amplitude randomness

So far we have used PWC input functions with normally distributed amplitudes (Fig. 3-a) motivated by the fact that in the limit of Δ → 0, *I_η_* approaches white noise. Earlier in the paper we have shown that the ability of PWC inputs to capture the resonant properties of a given cell depends on the multiple ways the receiving cell’s autonomous transient dynamics are activated by the different initial conditions resulting from the transition between different linear pieces, while randomness does not play any significant role in this process. Here we extend these ideas to the cells’ response variability. To this end, we use fully deterministic distributions of amplitudes within some range as described in Section 2.2.2 leaving the choice of the subset of permutations (number of trials) for each protocol as the only source of uncertainty in the input signal.

Figs. 9, S2 and S3 show that for the three types of deterministic distributions presented in Fig. 3, the PWC inputs uncover the voltage oscillatory properties of receiving cells (top panels) and show the same type of variability as the responses to the normally distributed PWC inputs discussed above (bottom panels). The differences are in the details. The main observed differences (by inspection) are in the responses to the P cells (panels a2) where the Var *P_η_* profiles are denser for the equispaced than for the bell-shaped-like PWCs and differences in the values of *D* for the latter are reflected in the peaks-and-troughs distributions in the Var*P_η_* profiles. A more detailed analysis is beyond the scope of this paper.

**Figure 9:**
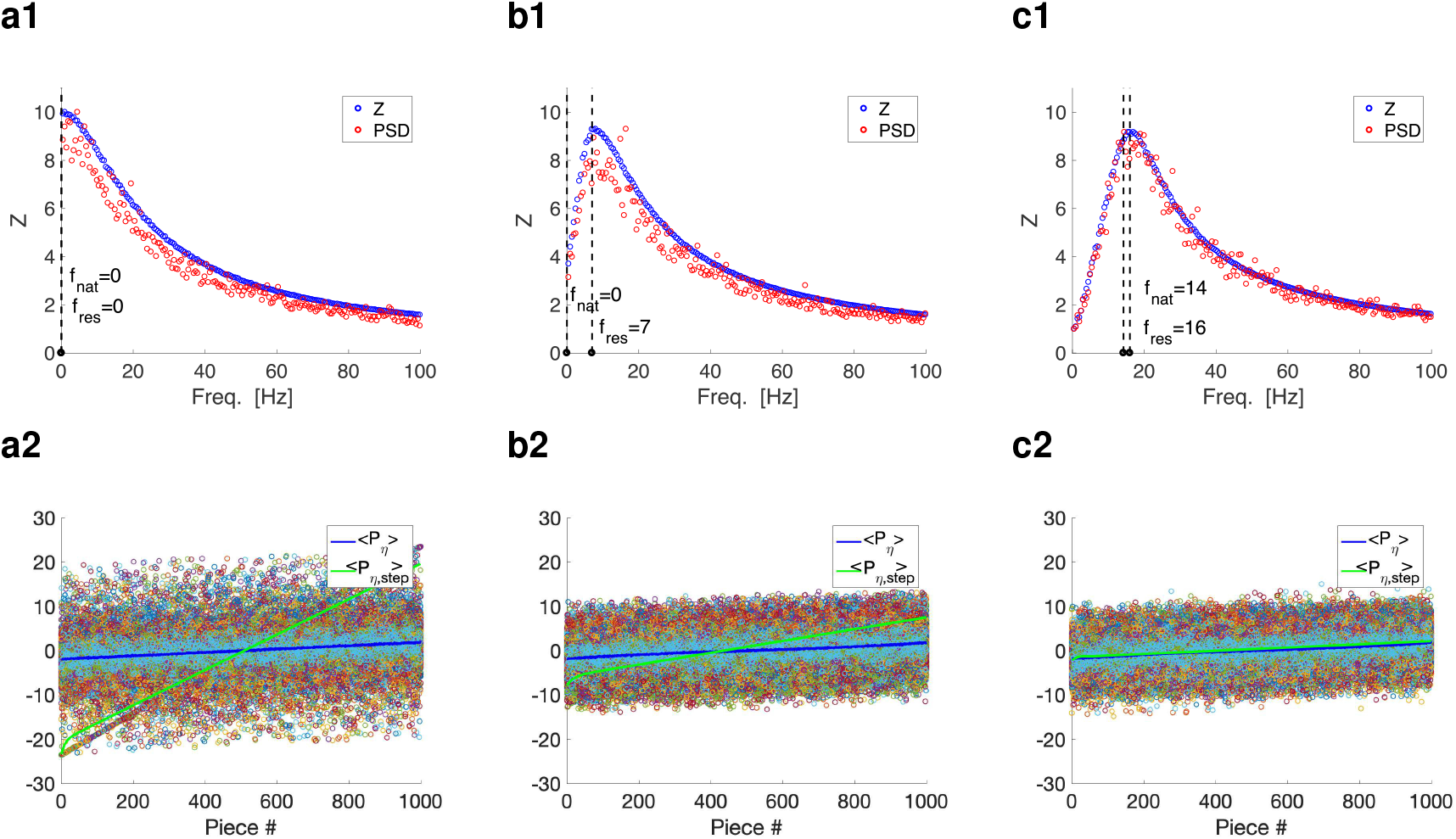
Piecewise constant inputs with arbitrarily ordered, but equispaced (non-randomly) distributed amplitudes capture the oscillatory properties of the target cells and the variability resulting from the autonomous transient dynamics. The piecewise constant inputs *I_η_* and *I_η,step_* have constant pieces with equispaced distributed amplitudes (equal amplitude differences) in the interval [−2, 2] with Δ = 1 (total time = 1000000 ms for row 1 and total time = 1000 ms for each trial for row 2. Similar results (with less resolution) are obtained for a smaller number of pieces for row 1 (total time = 10000 ms). Trials consist of different permutations (1000) of the same set of constant pieces 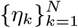. **a.** *g*_1_ = 0. **b.** *g*_1_ = 0.2. **c.** *g*_1_ = 1. We used the additional parameter values: *C* = 1, *g_L_* = 0.1 and *τ*_1_ = 100. **Top.** Impedance amplitude (*Z*) and (rescaled) power spectra density (PSD) profiles for the sample *V* trace for the responses to *I_η_* (random) and *I_η,step_* (ordered). The PSD profiles were rescaled so that the maxima of the PSD and *Z* profiles coincide. **Bottom**. The color dots indicate the peaks-and-troughs patterns for all trials reorganized so that the corresponding values of *η_k_* from which they originate are ordered in a monotonically increasing manner. All dots for a given piece correspond to the same value of *η_k_*. *< P_η_ >* (blue) is the mean value of the reordered peaks-and-troughs patterns for each linear piece. *< P_η,step_ >* (green) is the peaks-and-troughs pattern corresponding to the (ordered) input function *I_η,step_*.

### 3.5 The activation of the autonomous transient dynamics by the piecewise constant inputs with small Δ evokes the steady-state oscillatory properties of the receiving cells, and not their transient oscillatory properties

Along the previous sections, we have shown a number of examples where both the impedance amplitude *Z*(*f*) and the voltage PSD profiles of cells receiving PWC inputs with small durations Δ exhibit resonance, independently of whether the corresponding autonomous systems are a node (no intrinsic oscillations) or a focus (intrinsic damped oscillations). The impedance amplitude profile *Z*(*f*) of a system captures its steady-state response to oscillatory inputs. For linear systems, it is the magnitude of the complex-valued coefficient of the particular solution to the system (see Appendix A.2) forced by a sinusoidal function. In this calculation, the transient component of the solution to the forced system (the solution to the corresponding homogeneous system) is ignored. The resonance frequency *f_res_*, the peak frequency of *Z*(*f*), is different from the natural frequency *f_nat_* (the frequency of the transient damped oscillations). The latter is computed from the eigenvalues for the homogeneous system (see Appendices A.1 and A.2, and also [16, 21]), which controls the autonomous transient dynamics.

On the other hand, the dynamic mechanisms of generation of resonance involve the autonomous transient dynamics as uncovered by using dynamical systems tools [17, 18]. Most significantly, the interplay of the input frequency (whose inverse is a measure of the input time scale) and the cell’s intrinsic time scale determines the direction of motion of the response limit cycle trajectory in the phase-space. For the resonance frequency, this interaction of time scales produces the limit cycle trajectory with the maximal amplitude in the *v* direction. This mechanism does not rely on the cell’s eigenvalues and eigenvectors (autonomous steady-state dynamics) and it underlies the resonant responses of both N- and F-cells.

Because white noise has a constant PSD, both the *Z* and the *V* PSD profiles have the same frequency-dependent properties. This extends to PWC inputs with small enough values of Δ (e.g., Figs. 2a2 to -c2, Figs. 9- and S2- to S3-a1 to -b1), but not necessarily for large values of Δ (e.g., Figs. 2a2 to -c2, insets). Fig. 9-b1 (see also Figs. S2- to S3-b1) shows that both the *Z* and the *V* PSD profiles peak at *f_res_* when *f_nat_* = 0 (the receiving cell is a node). For these parameter values there is a clear separation between the cell’s response to the oscillatory input and the autonomous intrinsic dynamics (captured by *f_nat_*). In Fig. 9-c1 (see also Figs. S2-c1 to S3-c1) the *V* PSD profile seems to be superimposed to the *Z* profile, but because *f_res_* and *f_nat_* are very close, it is not clear whether the *V* PSD profile peaks at *f_res_*, *f_nat_* or in between. For the parameter values in Fig. 10-a1 the separation between *f_res_* and *f_nat_* is bigger and the *V* PSD profile clearly peaks around *f_res_* and far away from *f_nat_* for Δ = 1.

**Figure 10:**
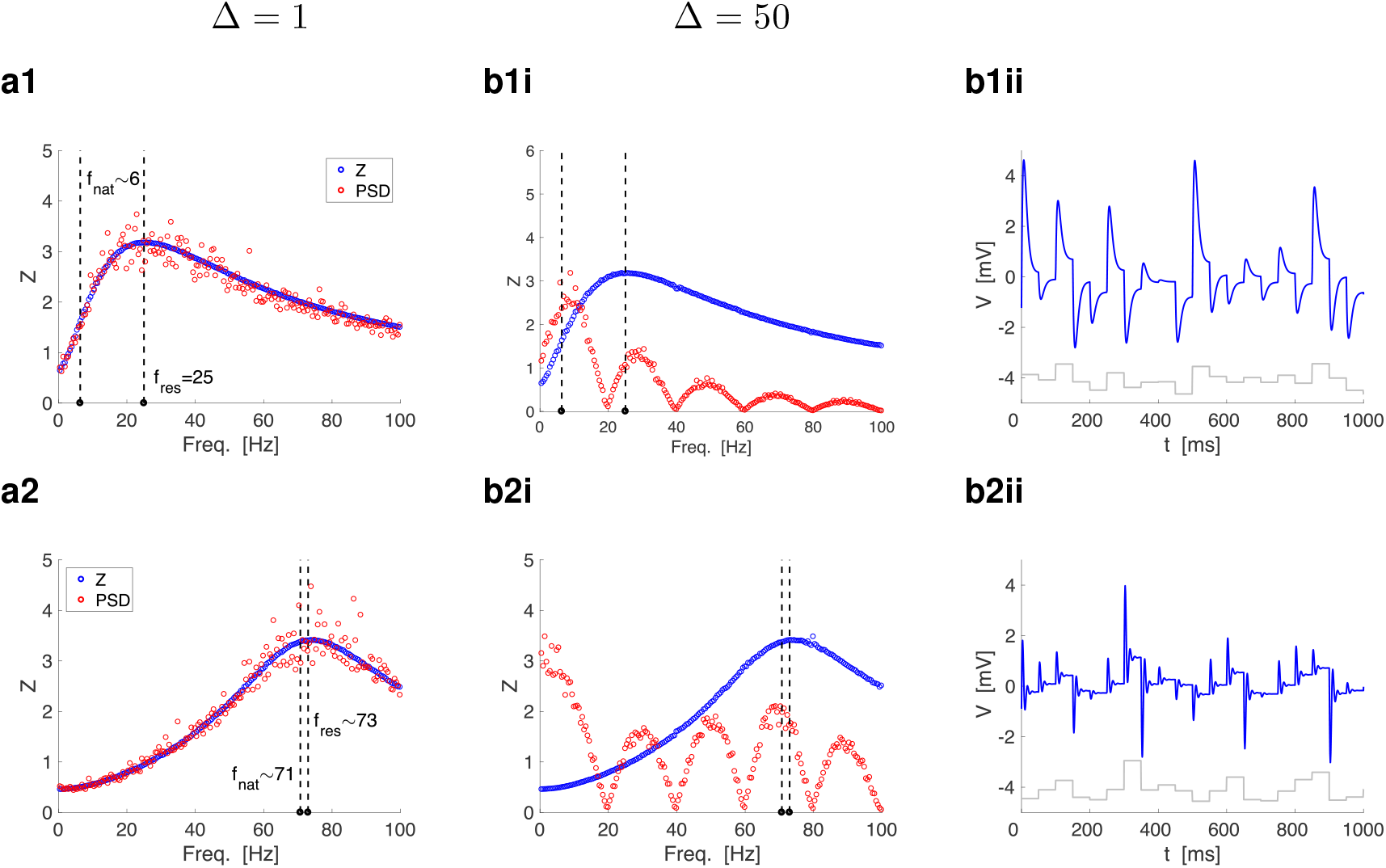
Cell’s responses to piecewise constant inputs with normally distributed amplitudes and different constant piece durations. The piecewise constant inputs *I_η_* have a total time equal to 1,000,000. **a.** Δ = 1. **b** Δ = 50. **Top.** 2D linear cell having a focus (F) and resonance with well separated values of *f_nat_* and *f_res_*. We used the following parameter values: *C* = 1, *g_L_* = 0.3, *g*_1_ = 1.3 and *τ*_1_ = 60. **Bottom.** 2D linear cells having a focus (F) and resonance with similar values of *f_nat_* and *f_res_* We used the following parameter values: *C* = 1, *g_L_* = 0.2, *g*_1_ = 2 and *τ*_1_ = 10. **Left and Middle.** Superimposed *Z* and voltage PSD profiles. The dashed vertical lines indicate *f_res_* (right) and *f_nat_* (left). **Right.** Sample voltage trace (blue). The gray curve is a caricature of *I_η_*.

In contrast, Figs. 10-b show more complex responses for Δ = 50. The duration of the constant pieces Δ determines an input time scale that interacts with the cell’s intrinsic time scale to produce the *V* response. This is true for all values of Δ, but for small values of Δ the corresponding frequencies (1000/Δ) are very large, away from the resonant frequency band, and the response time to each constant input piece very small. As Δ increases, the interaction between time scales is stronger and felt for the lower frequencies. For example, in Figs. 10-b for Δ = 50, the *V* PSDs show resonances occurring at multiples of 1000/Δ = 20. The shape of these resonant patterns reflects the properties of the autonomous transient dynamics rather than the properties of the corresponding *Z* profiles. In Figs. 10-b1, the sequence of PSD peaks decreases, while in Figs. 10-b2, the sequence of PSD peaks increases and the maximum of these peaks (save the maximum at *f* = 0) occurs roughly at *f_nat_*. This reflects the stronger oscillatory responses evoked by each constant piece due to their autonomous transient dynamics.

In summary, the autonomous transient dynamics plays a role in shaping the *V* response PSD patterns, but these patterns transition from reflecting the stationary properties of the *V* response to oscillatory inputs, captured by the *Z* profile, to the intrinsic properties of the receiving cell as Δ increases.

## 4 Discussion

Neuronal systems are subject to fluctuations either intrinsically or externally [23–32, 82, 83], which have been modeled as random Gaussian white or colored noise [39]. Cells subject to variable inputs have been shown to exhibit a number of non-expected behaviors (see Introduction for more details and references), including oscillatory voltage responses and resonances, and they also exhibit variability across trials. These phenomena result from the dynamic, often non-trivial interplay of the cell’s intrinsic properties and the properties of the noise. While variable inputs are ubiquitous, in particular in neuronal systems [24, 39, 84], several aspects of the dynamic mechanisms governing these interactions are not well understood. In particular, it is not well understood what aspects of the cell’s intrinsic dynamics play a role in each of these interactions, what properties of the variability are necessary, if at all, and how the two are integrated to produce these behaviors. Linear systems receiving Gaussian white noise can be solved analytically [75–77], but, in general, the resulting formulas provide limited mechanistic information. Nonlinear systems are generally not amenable to analytical solutions.

The investigation of dynamical systems primarily focuses on stationary solutions [85–87] and the properties of the transient behavior are less explored or even overlooked. Attractor networks are a key concept in neural computation [88–94] and have been proposed to capture a variety of cognitive process (e.g., working memory, navigation). Recently, transient dynamics have been argued to be important for neuronal processing and the encoding of information [95–101].

In previous work we showed that the cell’s autonomous transient dynamics (the cell’s transient voltage response to constant inputs, which is uncovered by the voltage response to abrupt changes between two constant inputs) play a significant role in the mechanistic explanation of the generation of resonance in linear and non-linear systems in response to oscillatory inputs [17, 18]. Even for linear systems for which the impedance amplitude and phase profiles can be computed analytically, the resulting formulas do not explain why and under what circumstances resonance emerges in systems that do not display intrinsic oscillations (i.e., systems having only real eigenvalues). We developed a dynamical systems approach to show that the cell’s response to oscillatory inputs can be thought of as a continuous sequence of transient voltage responses to constant inputs equal to the value of the input at a given time. The response trajectory for each input frequency tracks the cyclic motion of the voltage nullcline in the extended phase-plane diagram (technically, the projection of a surface in the three-dimensional space) and this evolution is governed by the properties of the autonomous transient dynamics. The resonant frequency corresponds to the optimal balance between the effective time scale at which the system transiently reacts to constant inputs and the input frequency. This could be captured by approximating the sinusoidal inputs by piecewise constant (PWC) functions with short-duration pieces and computing the sequence of (property joined) voltage response trajectories to these PWC inputs. Because in general these trajectories do not reach a close enough vicinity of the equilibrium for each constant piece, knowledge of the steady-state properties of the unforced target cell (eigenvalues and eigenvectors) does not inform the generation of resonance. We reasoned that similar ideas could shed light on the issues raised in the previous paragraph.

In this paper we set out to investigate the mechanisms of generation of oscillations and variability in response to PWC inputs *I_η_* with short-duration constant pieces and variable amplitudes. For each constant piece, the autonomous transient dynamics are able to develop, but the voltage response does not reach a close vicinity of the corresponding steady-state. For randomly distributed amplitudes *η*, the PWC inputs provide an approximation to Gaussian white noise [78]. However, although our results have implications for systems subject to random Gaussian noise, we are not making precise statements about the transition from the responses to the deterministic PWC inputs to the responses to stochastic Gaussian noise inputs as the duration of the constant pieces approaches zero. This requires more research and is beyond the scope of this paper.

We showed that oscillatory behavior can be generated in response to additive PWC inputs *I_η_* in both linear and nonlinear systems, independently of whether the stable equilibria of the unperturbed systems are foci or nodes. These results persisted when the amplitudes of the constant pieces were not randomly distributed, but chosen following a deterministic rule. For linear systems, this captures earlier results described in [76] for Gaussian white noise (see also [79] for two-dimensional linear cells with foci). However, the dynamic mechanisms of generation of oscillations and their dependence on the autonomous transient dynamics, in particular for cells having a node were not known. To understand the effects of the nonlinearities on the oscillatory voltage response we used a piecewise linear model (PWL) developed in [17] that mimics the voltage-dependences present in neuronal models in the voltage nullclines (for the current-balance equation). We showed that the oscillatory voltage response is amplified by the nonlinearities. This nonlinear amplification is consistent with previous work on resonance (for sinusoidal inputs) [17] and can be also explained in terms of the interaction between autonomous transient dynamics and the input properties. The PWL model has two regimes, the linear regime, in the vicinity of the fixed-point, and the nonlinear regime for voltage values higher than the “breaking point” determining the boundary between the two PWL input pieces (see Fig. 5-b). The nonlinear regime is in itself linear with respect to a virtual fixed-point, which is different from the actual fixed-point (the origin in Fig. 5-b). For each constant input piece for which the response trajectory crosses to the nonlinear regime, this trajectory is able to reach larger voltage values as compared to the linear model for the same input. This is because the slope of the voltage nullcline in the nonlinear regime is larger (less negative) than the slope in the linear regime, and therefore the trajectory for transient voltage response to the corresponding constant input is able to jump to higher values as compared to the linear regime. This clearly depends on the properties of the PWL model, in particular on the slope of the “broken” linear piece, which we chose to be larger (less negative) than the linear piece in the linear regime (see Fig. 5-b). Other slopes may lead to attenuation of the voltage response. Following similar ideas, we showed that the voltage response to multiplicative conductance-based synaptic inputs is attenuated as compared to the underlying model with additive noise. In this case, the “broken” piece is more negative than the linear one (see Fig. 5-c). This has mechanistic implications for the experimental results discussed in [102], comparing the voltage responses of current- and conductance-based synaptic fluctuations. A more detailed explanation requires the use of high-frequency Poisson-distributed synaptic-like inputs. We analyze this in more detail in a companion paper [103].

The effect of the autonomous transient dynamics on the response variability to *I_η_* is already apparent in the simple illustrative example in Figs. 1-c to -e. The voltage response variability emerges as the result of the transition between regimes corresponding to different constant pieces. The variability is minimal for the response to *I_η,step_* (Figs. 1-c1 to -e1) where the constant pieces are arranged by increasing amplitude and the variability increases as the constant pieces are permuted (Figs. 1-c2 to -e2) since the transient voltage response for each constant piece regime depends on the initial conditions for that regime. These, in turn, depend on the value of the participating variables at the end of the previous regime, which is different for different trials (different permutations of the order of the constant pieces). To clarify these ideas we designed protocols where the set of amplitudes *η* was the same for all trials and each trial used a different permutation of this set. We then rearranged the voltage response peak-and-trough profiles *P_η_* in increasing order of input amplitudes *η*. In this way, we were able to compare the different voltage responses to the same input amplitude across trials. We used both randomly distributed amplitudes and deterministic distribution of amplitudes. For each input amplitude *η_k_*, the variability in *P_η_k* across trials was the result of the multiple different ways the target cell reacts to the input *η_k_*. Again, this is due to the differences in the initial conditions for that regime, which in turn depend on the values of the participating variables at the end of the previous regime. This variability was strongly affected by the properties of the target cells and their ability to produce overshoots and damped oscillations in response to abrupt changes in constant inputs, which in turn depends on the model parameters, which constrain these responses. This interpretation is strengthened by the fact that the variability patterns remained qualitatively the same when we relaxed the requirement that the amplitudes are randomly distributed and we used permutations of the same deterministic distribution for each protocol. Although it was not the full overshoot or the damped oscillations that affected the voltage response, but rather the initial portion to these responses, the particular dependence on the model parameters remains. In other words, cells that exhibit stronger autonomous transient dynamics show more variability. In this sense, this is consistent with the results for one-dimensional OU processes and it is intuitive for higher-dimensional OU processes, however, it is not directly clear from the complex covariance formulas. For nonlinear processes, analytic computations are not possible and therefore the method we developed allows to make predictions based on the knowledge of the autonomous transient dynamics and the structure of the input. Our results further predict that the variability would be reduced if the input functions involve gradual rather than abrupt transitions.

These predictions and the predictions about the oscillatory properties of cells in response to PWC inputs could be tested experimentally *in vitro* using current clamp techniques or *in vivo* using optogenetic tools [104–106]. In the first case, variability in response to PWC inputs is expected to be well correlated with appropriate metrics for the autonomous intrinsic dynamics measured in response to constant inputs. Obtaining these correlations in the second case could be more challenging, but still possible by using the appropriate tools to measure the response membrane potentials.

Time-dependent inputs can be approximated by a discrete sequence of short-duration constant pieces (e.g., Fig. 1-a) and therefore the sequence of the properly joined transient solutions to these constant pieces (Figs. 1-c to -e) provides an approximation to the system’s voltage response to the original (time-dependent) inputs. The short duration of the constant pieces does not allow for the corresponding steady-state solutions to develop and therefore the information about the input is encoded by the transient trajectories. Complex odors (different types of stimuli at different concentrations) and their representations have been proposed to operate in a similar way [95, 101]. A potentially more involved experimental test or application of the ideas presented in this paper is the presentation of complex odors consisting of sequences of different permutations of a number of “basic odors” [95, 101]. Variability in the responses is expected across trials and an increase in the variability patterns is expected as the “intensity” of these odors increase. However, odor representations in the olfactory research is a network phenomenon and therefore additional research is needed to extend our results to include network effects.

Our results have also implications for the understanding of the emergence of variability in neuronal systems [107] and dynamical systems in general, and in the absence of noise. In the simple systems we used here, variability emerges as the result of the interaction between the system’s autonomous transient dynamics and a set of deterministic inputs where the only source of uncertainty is the choice of the subset of all possible PWC functions with the same set of constant piece amplitudes. Key for this variability to emerge are the abrupt transitions between constant piece regimes and the sensitivity to initial conditions of the autonomous transient dynamics. In higher-order networks, this can be evoked by the network internal dynamics without the need of external inputs. Therefore our results have implications for the encoding of information in general and for these resulting from expansions or quenching of variability [3, 108]. More research is needed to establish in these networks how variability is affected by the node and connectivity parameters. Finally, if variability encodes the autonomous transient dynamics of the target cells, then decoding methods should be able to infer these dynamics. These also require more research.

## Acknowledgments

This work was partially supported by the National Science Foundation grant DMS-1608077 (HGR).

## Ethics declarations

**Conflict of interest:** The authors declare that they have no conflict of interest.

## A Intrinsic and resonant oscillatory properties of 2D linear systems

Consider

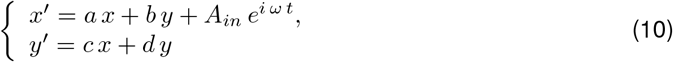

where *a*, *b*, *c* and *d* are constants, *ω* = 2*πf/*1000 > 0 is the input frequency and *A_in_* ≥ 0 is the input amplitude. The prime sign represents the derivative with respect to *t*. The units of *t* are ms and the units of *f* are Hz.

## A.1 Intrinsic oscillations

The characteristic polynomial for the corresponding homogeneous system (*A_in_* = 0) is given by

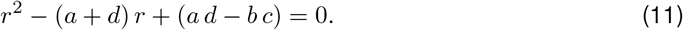

The eigenvalues are given by

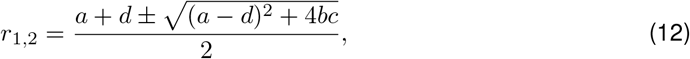

and the natural (intrinsic) frequency of the (damped) oscillations (in Hz if *t* has units of ms) is given by

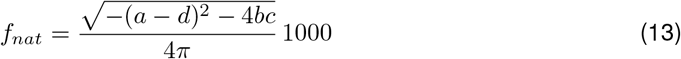

assuming (*a* − *d*)^2^ + 4*bc* < 0.

## A.2 Resonance and the impedance amplitude profile

The impedance amplitude profile *Z*(*ω*) for system (10)-(11) is the magnitude

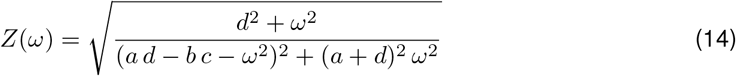

of the complex valued coefficient of the particular solution to the system

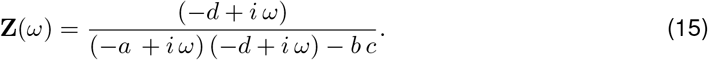

For 1D system, these quantities are given, respectively, by

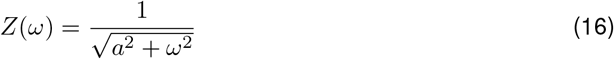

and

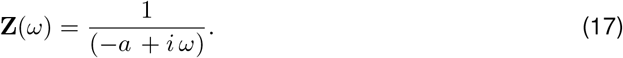

The resonance frequency *f_res_* (in Hz if *t* has units of ms) is the frequency at which *Z* reaches its maximum

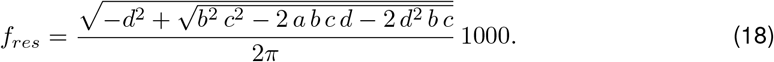

## A.3 Response to constant inputs

The equilibrium solution to system (10) for a constant input *A_in_* (i.e., *ω* = 0) is given by

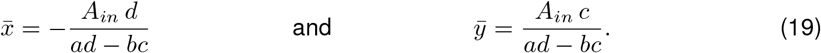

The eigenvectors are given by

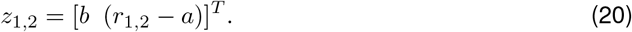

The solution satisfying the initial conditions [*x*(0) *y*(0)]^*T*^ = [*x*_0_ *y*_0_]^*T*^ is given by

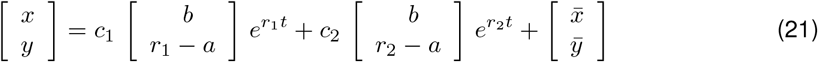

where

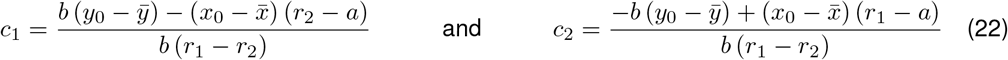

For 1D systems (b = 0),

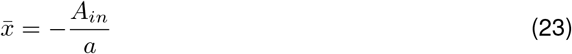

and

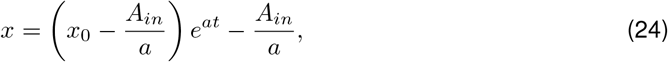

where *x*(0) = *x*_0_

## B Ornstein-Uhlenbeck (OU) Process

## B.1 One-dimensional OU process

The 1D OU process [75] is described by the following linear stochastic differential equation

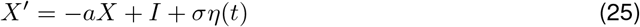

where *a >* 0, *I* and *σ* are constants and *η*(*t*) is zero-mean and *δ*-correlated Gaussian white noise. The parameter *a* is the inverse of the time constant and measure the strength by which the system reacts to perturbations. The parameter *σ* measures the intensity of the noise. The quotient *I/α* is the asymptotic mean.

Using standard methods [76, 77] one can compute the solution satisfying *X*(0) = *x*_0_, which is the sum of a deterministic function with the form (24) and an integral of a deterministic function with respect to a Wiener process. The solution is normally distributed with mean and variance given, respectively by

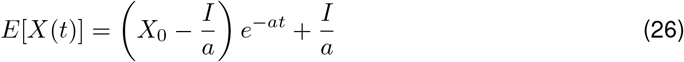

and

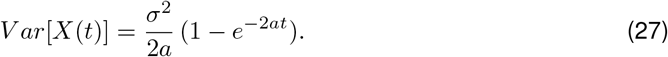

## B.2 Higher-dimensional OU process

The multivariate OU process [75] is described by the following linear stochastic differential equation

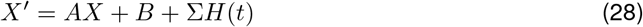

where *X* is an *n*-dimensional vector, *A* is an *n × n* matrix, *B* is an *n*-dimensional vector, Σ is an *n × m* matrix and *H* is a vector of independent zero-mean and *δ*-correlated Gaussian white noise components. Using standard methods [76, 77] one can compute the solution satisfying *X*(0) = *x*_0_.

The solution is normally distributed. The mean is given by

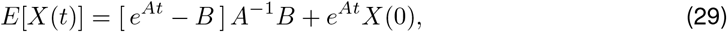

and the covariance matrix is given by

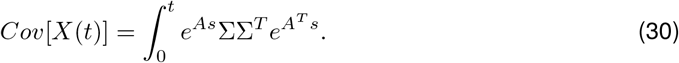

Under certain conditions, the covariance matrix corresponding to the stationary solutions reads

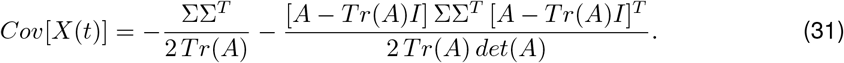

## Supplementary Material

**Figure S1:**
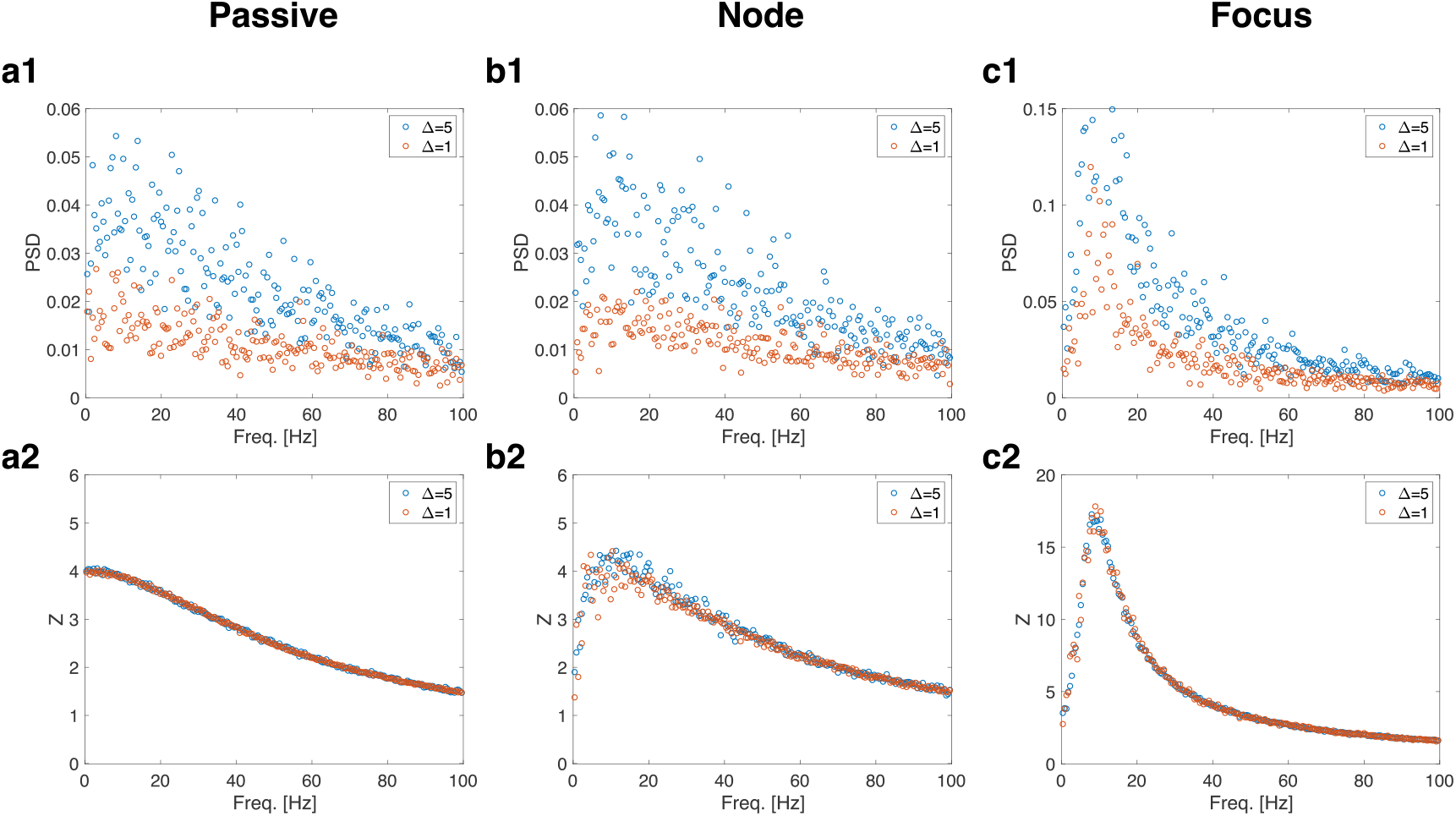
Piecewise constant inputs with arbitrarily ordered, but uniformly (non-randomly) distributed amplitudes capture the transient dynamics of the target cells. The piecewise constant inputs *I_η_* have Δ = 5 (total time = 10000 ms, N = 2000). The parameter values are the same as in Figs. 1 and 2. **Row 1.** Power spectra density (PSD) profiles for the sample *V* trace. **Row 2.** Impedance amplitude (*Z*) profiles for the sample *V* trace. **a.** Passive cell (*f_nat_* = *f_res_* = 0). We used the following paramete.r values: *C* = 1, *g_L_* = 0.25. **b.** 2D linear system exhibiting an overshoot in response to step-constant inputs ((*f_nat_* = 0*, f_res_* ~ 9*Hz*). We used the following parameter values: *C* = 1, *g_L_* = 0.25, *g*_1_ = 0.25, *τ*_1_ = 100. **c.** 2D linear system exhibiting damped oscillations in response to step-constant inputs (*f_nat_* ~ *f_res_* ~ 8*Hz*). We used the following parameter values: *C* = 1, *g_L_* = 0.05, *g*_1_ = 0.3, *τ*_1_ = 100. Notice different values of Δ as shown in the legends.

**Figure S2:**
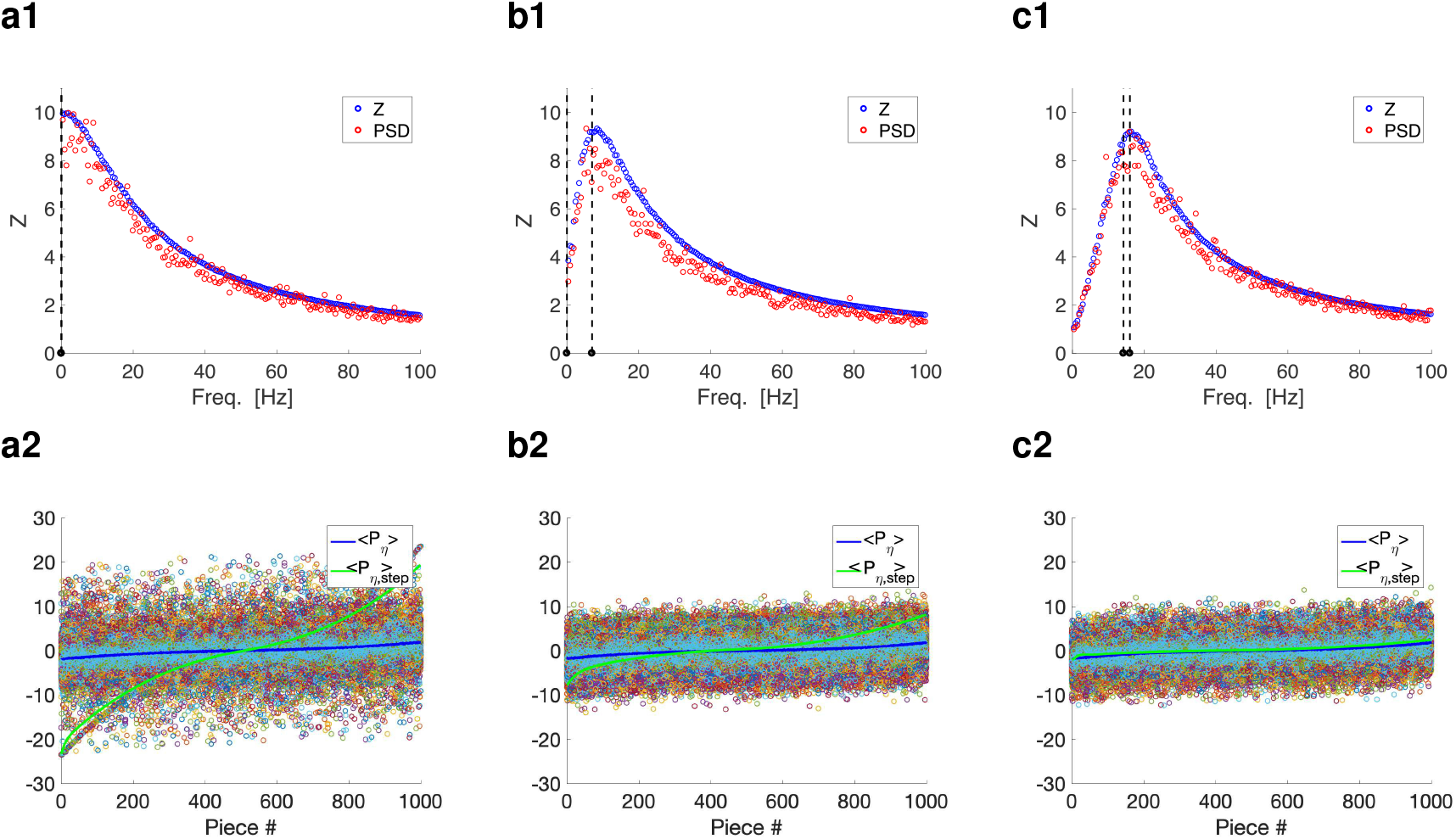
Piecewise constant inputs with arbitrarily ordered, but bell-shaped (non-randomly) distributed amplitudes capture the oscillatory properties of the target cells and the variability resulting from the autonomous transient dynamics. The piecewise constant inputs *I_η_* and *I_η,step_* have constant pieces with (non-random) bell shapped distributed amplitudes in the interval [−2, 2] with Δ = 1 (total time = 1000000 ms for row 1 and total time = 1000 ms for each trial for row 2. Similar results (with less resolution, variance = 1.5) are obtained for a smaller number of pieces for row 1 (total time = 10000 ms). Trials consist of different permutations (1000) of the same set of constant pieces 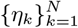. **a.** *g*_1_ = 0. **b.** *g*_1_ = 0.2. **c.** *g*_1_ = 1. We used the additional parameter values: *C* = 1, *g_L_* = 0.1 and *τ*_1_ = 100. **Top.** Impedance amplitude (*Z*) and (rescaled) power spectra density (PSD) profiles for the sample *V* trace for the responses to *I_η_* (random) and *I_η,step_* (ordered). The PSD profiles were rescaled so that the maxima of the PSD and *Z* profiles coincide. **Bottom**. The color dots indicate the peaks-and-troughs patterns for all trials reorganized so that the corresponding values of *η_k_* from which they originate are ordered in a monotonically increasing manner. All dots for a given piece correspond to the same value of *η_k_*. *< P_η_ >* (blue) is the mean value of the reordered peaks-and-troughs patterns for each linear piece. *< P_η,step_ >* (green) is the peaks-and-troughs pattern corresponding to the (ordered) input function *I_η,step_*.

**Figure S3:**
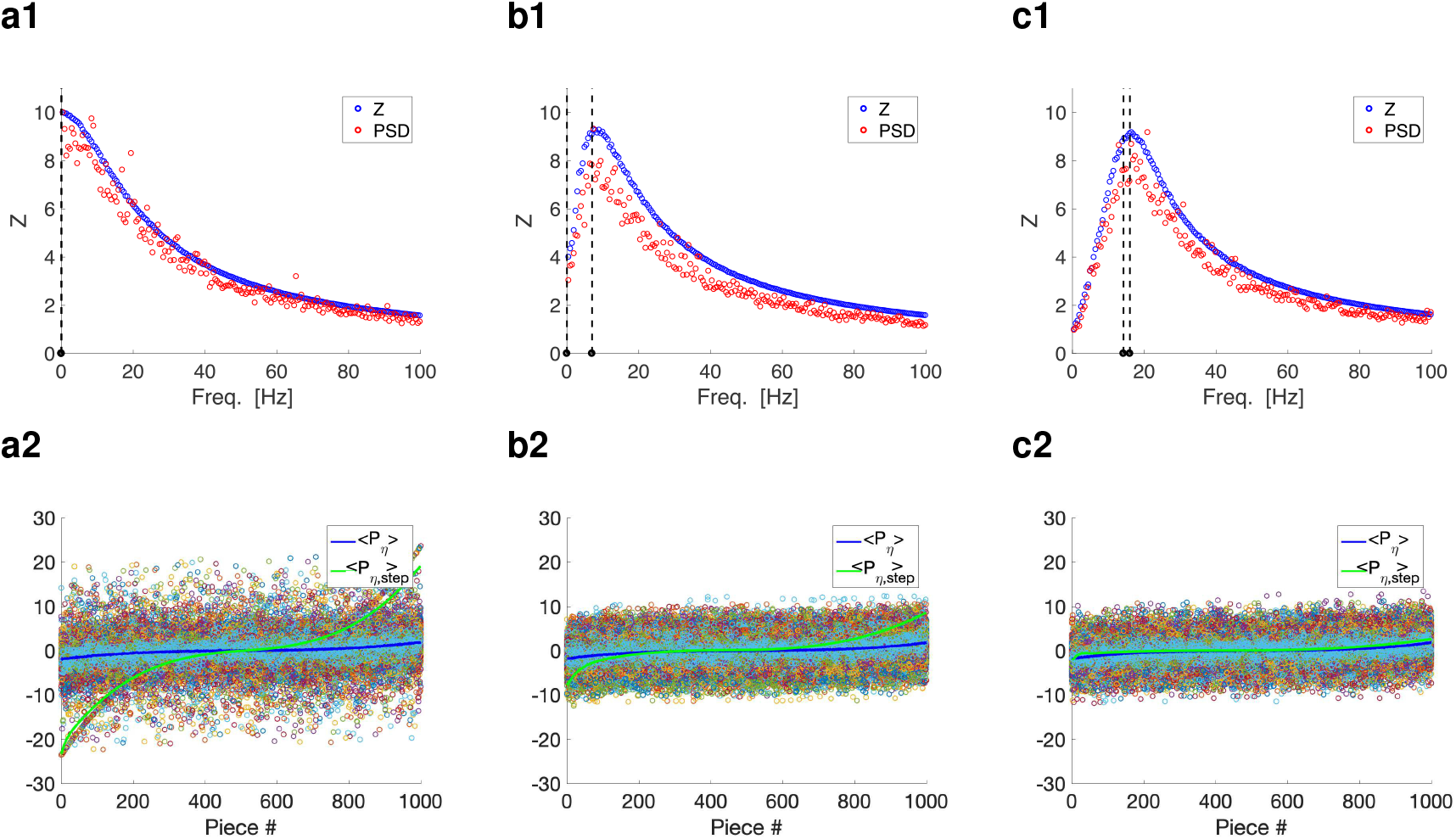
Piecewise constant inputs with arbitrarily ordered, but bell-shaped (non-randomly) distributed amplitudes capture the oscillatory properties of the target cells and the variability resulting from the autonomous transient dynamics. The piecewise constant inputs *I_η_* and *I_η,step_* have constant pieces with (non-random) bell shapped distributed amplitudes in the interval [−2, 2] with Δ = 1 (total time = 1000000 ms for row 1 and total time = 1000 ms for each trial for row 2. Similar results (with less resolution, variance = 1) are obtained for a smaller number of pieces for row 1 (total time = 10000 ms). Trials consist of different permutations (1000) of the same set of constant pieces 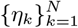. **a.** *g*_1_ = 0. **b.** *g*_1_ = 0.2. **c.***g*_1_ = 1. We used the additional parameter values: *C* = 1, *g_L_* = 0.1 and *τ*_1_ = 100. **Top.** Impedance amplitude (*Z*) and (rescaled) power spectra density (PSD) profiles for the sample *V* trace for the responses to *I_η_* (random) and *I_η,step_* (ordered). The PSD profiles were rescaled so that the maxima of the PSD and *Z* profiles coincide. **Bottom**. The color dots indicate the peaks-and-troughs patterns for all trials reorganized so that the corresponding values of *η_k_* from which they originate are ordered in a monotonically increasing manner. All dots for a given piece correspond to the same value of *η_k_*. *< P_η_ >* (blue) is the mean value of the reordered peaks-and-troughs patterns for each linear piece. *< P_η,step_ >* (green) is the peaks-and-troughs pattern corresponding to the (ordered) input function *I_η,step_*.

